# The *C. elegans* immune switch proteins PALS-25 and PALS-22 localize to mitochondria and regulate fragmentation

**DOI:** 10.1101/2025.10.13.682198

**Authors:** Spencer S. Gang, Max W. Strul, Emily R. Troemel

## Abstract

The nematode *C. elegans* controls immunity against intracellular pathogens like microsporidia using the *pals* gene family, which has expanded in *C. elegans* compared to mammals. *pals-22* is a negative regulator that restrains *pals-25*, which serves as a positive regulator of immunity. *pals-22* and *pals-25* encode proteins that bind each other and can act in the intestine and epidermis, but their subcellular localization and mechanism of action have not been described. Here we show that PALS-22 and PALS-25 proteins localize to mitochondria, with PALS-25 being required for PALS-22 localization to mitochondria. The C-terminus of PALS-25 is both necessary and sufficient for mitochondrial localization. Loss of PALS-22 causes mitochondrial fragmentation, which occurs after activating the Intracellular Pathogen Response (IPR), a transcriptional program induced by intracellular infection. Mitochondrial fragmentation induced in an independent manner increases resistance against microsporidia infection. Thus, PALS-22/25-mediated fragmentation of mitochondria appears to increase immunity against intracellular infection.

## Introduction

Microsporidia are ubiquitous fungal pathogens that infect almost all animal phyla and complete their entire replicative lifecycle inside host cells, similar to viruses^1,2^. In *C. elegans*, the transcriptional Intracellular Pathogen Response (IPR) provides defense against the natural microsporidian species *Nematocida parisii* and the Orsay virus, a single-stranded RNA virus^3^. While both pathogens trigger the IPR, only the Orsay virus appears to activate the IPR via the RIG-I homolog DRH-1 sensor^4,5^. Although *C. elegans* lacks classical interferons, this RIG-I-mediated activation of the IPR resembles the type-I interferon response (IFN-I response), a key anti-viral defense program in humans that also helps resist against microsporidia infections^6,7^.

Many IPR genes belong to the *pals* gene family, characterized by a divergent ‘*pals’* sequence signature found in *C. elegans,* mice and humans^8^. Humans and mice each have one *pals* gene ortholog, while *C. elegans* has at least 39 *pals* genes, divided into two classes: a) 26 infection-induced, and b) 13 non-induced. Among the 26 induced genes is *pals-14*, which promotes resistance to microsporidia^9^, and *pals-5*, which is commonly used as a reporter for IPR induction and can reduce levels of protein aggregates^10^. Some non-induced *pals* genes are negative regulators of the IPR (*pals-17* and *pals-22)*, while some others are positive regulators (*pals-25, pals-16* and *pals-20*)^11,12^. These IPR regulators act in ‘modules’, with *pals-17* repressing *pals-20/pals-16,* and *pals-22* repressing *pals-25*.

The best characterized module is composed of *pals-22* and *pals-25,* which act as an OFF/ON switch of the IPR. *pals-22* mutants have constitutive IPR gene expression, increased resistance to intestinal intracellular pathogens like microsporidia and virus, as well as increased silencing of repetitive genomic elements. All of these phenotypes are reversed in *pals-22 pals-25* double mutants^11^. Notably, *pals-22* and *pals-25* have also been shown to regulate immunity against oomycetes, which are natural eukaryotic pathogens of the epidermis^13^. *pals-22* mutants have slowed development and shortened lifespan, which is reversed by mutations in *pals-25*, indicating that PALS-22 and PALS-25 control a physiological switch from a growth state to a defense state^11^. PALS-22 and PALS-25 proteins bind to each other, and when this association is lost, PALS-25 can activate IPR genes and resistance to infection^14,15^. PALS-22 and PALS-25 are expressed broadly in *C. elegans*, with known roles in the intestine and epidermis^11,15,16^.

While the phenotypes, tissue expression, and protein-protein interactions of PALS-22 and PALS-25 have been described, their mechanism of action is still mysterious. Here, we show that both proteins localize to the mitochondria. PALS-22 depends on PALS-25 for its mitochondrial association, while PALS-25 localizes to mitochondria independently of PALS-22. Upon loss of PALS-22, which triggers an IPR activated state, PALS-25 protein forms puncta along the mitochondria. We find that the C-terminus of PALS-25 is necessary and sufficient for mitochondrial localization, and the N-terminal 40 amino acids of PALS-25 are required for its induction of the IPR. We also show that loss of PALS-22 leads to fragmentation of mitochondria, with a loss of branch number and shorter branch length. Kinetic studies of PALS-22 depletion, together with analysis of PALS-17 knockdown, suggest that IPR induction may trigger mitochondrial fragmentation. We also show that *N. parisii* associates with *C. elegans* mitochondria. Furthermore, when mitochondria are fragmented independent of IPR induction, there is increased resistance to *N. parisii* infection. Altogether, these studies indicate that PALS-22 and PALS-25 localize to mitochondria, and the induction of mitochondrial fragmentation helps promote resistance to microsporidia infection.

## Results

### PALS-22 and PALS-25 proteins localize to mitochondria

To investigate the subcellular localization of PALS-22 and PALS-25 proteins, we used TransgeneOme fosmids, which are a resource collection of specific *C. elegans* genes tagged at their C termini with GFP, surrounded by approximately 20 kb of endogenous regulatory region (Supp Fig 1A, B)^17^. Of note, *pals-22* and *pals-25* are found in an operon together (Supp Fig 1C). As previously described for strains carrying these TransgeneOme constructs^15^, PALS-22::GFP and PALS-25::GFP are expressed in many tissues in *C. elegans* including the intestine and epidermis. Using these strains, we focused on their subcellular expression pattern in the epidermis, because have strong expression in that tissue, particularly in seam cells. Furthermore, they can act in the epidermis to regulate the IPR^15^. Here, we observed that both PALS-22::GFP and PALS-25::GFP exhibit long, branched-like expression patterns that seemed characteristic of mitochondrial morphology. Indeed, when animals expressing PALS-22::GFP or PALS-25::GFP were stained with MitoTracker Red, we saw nearly perfect co-localization of both PALS-22::GFP and PALS-25::GFP with MitoTracker Red staining in the epidermis (Fig 1A, B). Imaging in the intestine can be confounded by autofluorescence, and PALS-25 expression was too low to assess its subcellular localization in this tissue. However, we did observe colocalization of PALS-22::GFP with MitoTracker Red in the intestine, similar to its localization in the epidermis (Supp Fig 1D).

**Figure 1.**
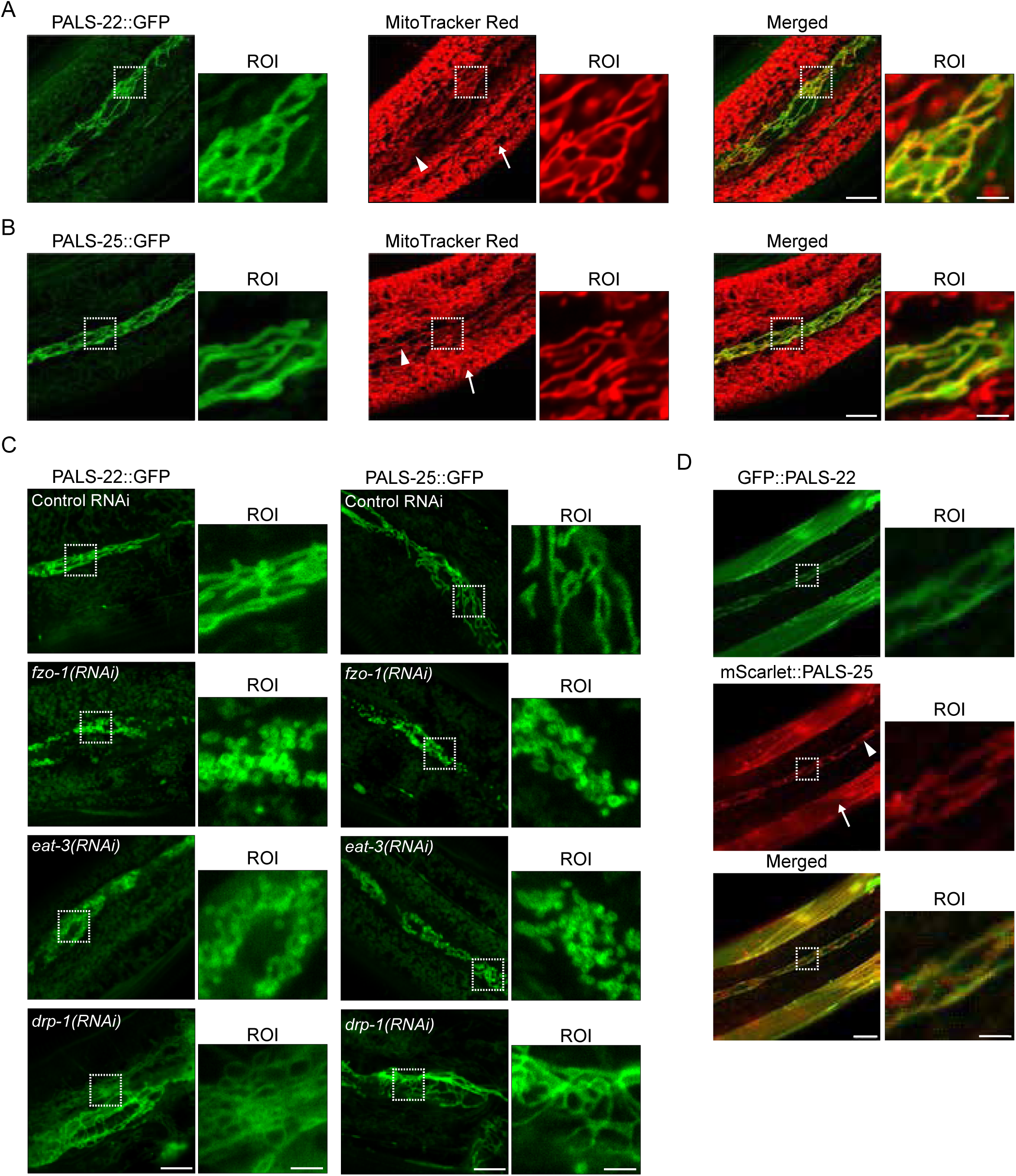
PALS-22 and PALS-25 are mitochondrially associated proteins. **A-B)** PALS-22::GFP **(A)** and PALS-25::GFP **(B)** colocalize with the mitochondria marker MitoTracker Red with robust expression in seam cells. The epidermis of adult animals is shown. White triangle denotes seam cell mitochondria, white arrow denotes non-seam cell epidermal mitochondria. **C)** Representative images of the epidermis of PALS-22::GFP (left) or PALS-25::GFP (right) adult animals following treatment with control, *fzo-1*, *eat-3*, and *drp-1* RNAi. **D)** GFP::PALS-22 and mScarlet::PALS-25 co-localize in muscle cells and epidermal cells in young adult animals. White triangle denotes seam cell mitochondria, white arrow denotes muscle. For **A-D**, scale bar = 10 μm, region of interest (ROI) scale bar = 2.5 μm.

We then performed further analysis of PALS-22 and PALS-25 localization to mitochondria. First, we performed RNAi against known fission and fusion factors that regulate mitochondria morphology^18^. Consistent with PALS-22 and PALS-25 localizing to mitochondria, we found that RNAi against the fusion factors *fzo-1* and *eat-3* led to more condensed PALS-22::GFP and PALS-25::GFP localization patterns resembling fragmented mitochondria (Fig 1C). We also found that RNAi against *drp-1,* whose wild-type function promotes fission of the outer mitochondrial membrane, led to longer network-like PALS-22::GFP and PALS-25::GFP localization patterns, typical of elongated mitochondria (Fig 1C). Next, to analyze PALS-22 and PALS-25 co-localization in the same cells, we generated a transgene that has PALS-22 tagged with GFP and PALS-25 tagged with the red fluorescent protein mScarlet. Here we made a ‘mini-operon’ of *pals-22* and *pals-25* cDNAs driven by 2 kb of *pals-22* upstream region, with fluorescent tags added to the N-termini of each gene (Supp Fig 1E). Again, we saw localization of both GFP::PALS-22 and mScarlet::PALS-25 to mitochondria-like structures in the epidermis, with strong co-localization of these two proteins in seam cells (Fig 1D).

Therefore, we conclude that PALS-22 and PALS-25 both localize to mitochondria and appear to be at the outer mitochondrial membrane instead of directly localizing with MitoTracker staining in the mitochondrial matrix (Fig 1 A, B and Supp Fig 1D). Further supporting the conclusion that PALS-22 and PALS-25 localize to the outer mitochondrial membrane is the observation that *drp-1* RNAi caused these two proteins to adopt a more elongated localization pattern (Fig 1C). Notably, *drp-1* is required for fission of the outer mitochondrial membrane, but not the inner mitochondrial membrane, and loss of *drp-1* will cause more elongated outer mitochondrial membranes, but fragmented inner mitochondrial membrane and matrices (^19,20^ and see below).

To confirm that these tagged proteins are functional, we crossed each of the TransgeneOme constructs into a *pals-22 pals-25(jy80)* strain background, which is a clean deletion of the *pals-22/pals-25* operon, and then analyzed IPR induction (Supp Fig 1C). Here we found that strains carrying either PALS-22::GFP or PALS-25::GFP were capable of inducing IPR gene expression upon RNAi knock-down of *pals-22* (Supp Fig 2A, B). Because disruption of the interaction between PALS-22 and PALS-25 can activate the IPR, we also analyzed basal IPR gene expression in these strains as a read-out for whether the tags might disrupt their interaction. Here, we did not see a significant induction of IPR gene expression (Supp Fig 2C). Therefore, the GFP tags do not appear to interfere with PALS-22 and PALS-25 normal interactions or with their ability to induce the IPR, and thus, we conclude that the PALS-22::GFP and PALS-25::GFP transgenes are functional.

### PALS-22 mitochondrial localization depends on PALS-25, which uses its C-terminus for mitochondrial localization

To examine how localization of PALS-22 and PALS-25 are regulated, we performed RNAi knock-down of *pals-25* and examined PALS-22::GFP localization. Here, we found that PALS-22::GFP lost its mitochondrial association, showing a diffuse cytosolic expression pattern instead (Fig 2A, Supp Fig 3A). When we performed Western blots, there was no significant decrease in PALS-22::GFP overall protein levels upon *pals-25* RNAi (Supp Fig 3B). In contrast, we found with Western blot analysis that there was a significant decrease in PALS-25::GFP protein levels upon RNAi knock-down of *pals-22*, which activates the IPR (Supp Fig 3C). This decrease in PALS-25 protein caused by *pals-22* RNAi cannot be explained by a decrease in *pals-25* mRNA (Supp Fig 2B). Furthermore, we found that PALS-25::GFP formed discrete puncta after *pals-22* RNAi, with these puncta still apparently localizing to mitochondria (Fig 2B, Supp Fig 3D). Therefore, PALS-22 depends on PALS-25 for its localization to mitochondria (Fig 2A). In contrast, PALS-25 does not require PALS-22 for its localization to mitochondria, but does require PALS-22 to maintain wild-type protein levels, and to maintain an even distribution pattern along these mitochondria (Fig 2B).

**Figure 2.**
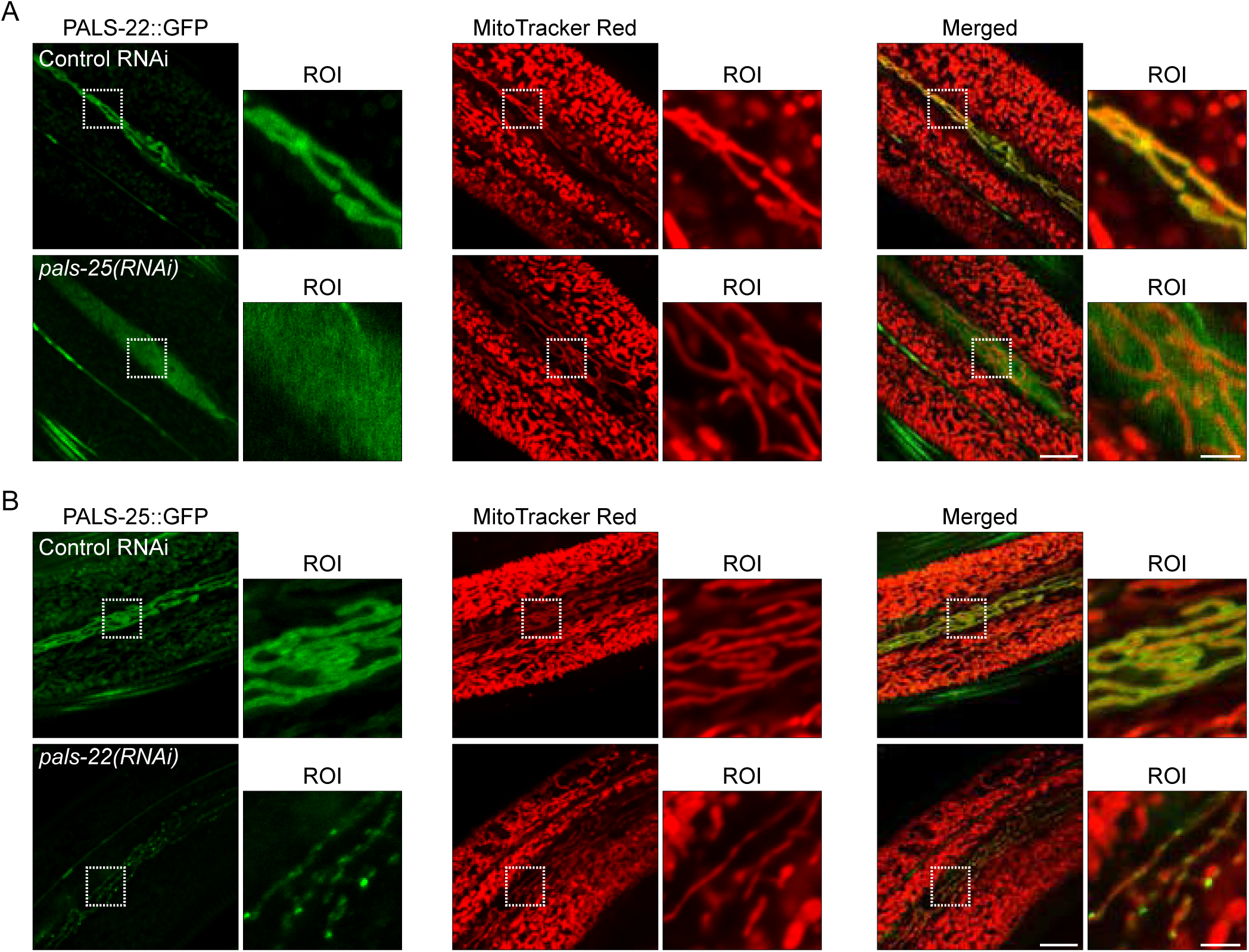
The dependence of PALS-22 and PALS-25 localization on each other. **A)** PALS-22::GFP expression and localization in the epidermis relative to mitochondria labeled with MitoTracker Red following treatment with control or *pals-25* RNAi. *pals-25* RNAi causes PALS-22::GFP to lose association with mitochondria in seam cells of the epidermis. **B)** PALS-25::GFP expression and localization in the epidermis relative to mitochondria labeled with MitoTracker Red following treatment with control or *pals-22* RNAi. *pals-22* RNAi, which induces the IPR, causes PALS-25 to form puncta localized to mitochondria in epidermal seam cells. For **A** and **B**, scale bar = 10 μm, ROI scale bar = 2.5 μm.

Given that PALS-25 appears to be the driver of PALS-22 localization to mitochondria, we next examined which domain of PALS-25 is responsible for mitochondrial localization. Our prior work determined that the last 13 amino acids at the C-terminus of PALS-25 are required for its association with PALS-22. This conclusion is based on analysis of the PALS-25 Q293* mutant, hereafter referred to as GOF (Gain Of Function), which has lost association with PALS-22, and thus has constitutive IPR gene expression (Supp Fig 1C, Fig 3A). Here, we examined whether PALS-25(GOF) still localizes to epidermal mitochondria through co-localization of mScarlet::PALS-25(GOF) and VWA-8::GFP, which localizes to mitochondria. Because of mScarlet aggregation likely related to the IPR activation in this strain, it was difficult to clearly co-localize mScarlet::PALS-25(GOF) to mitochondria. However, this protein did have a discontinuous localization pattern, similar to mitochondria in some areas, suggesting the C-terminal 13 amino acids may not be required for mitochondrial localization (Fig 3B).

**Figure 3.**
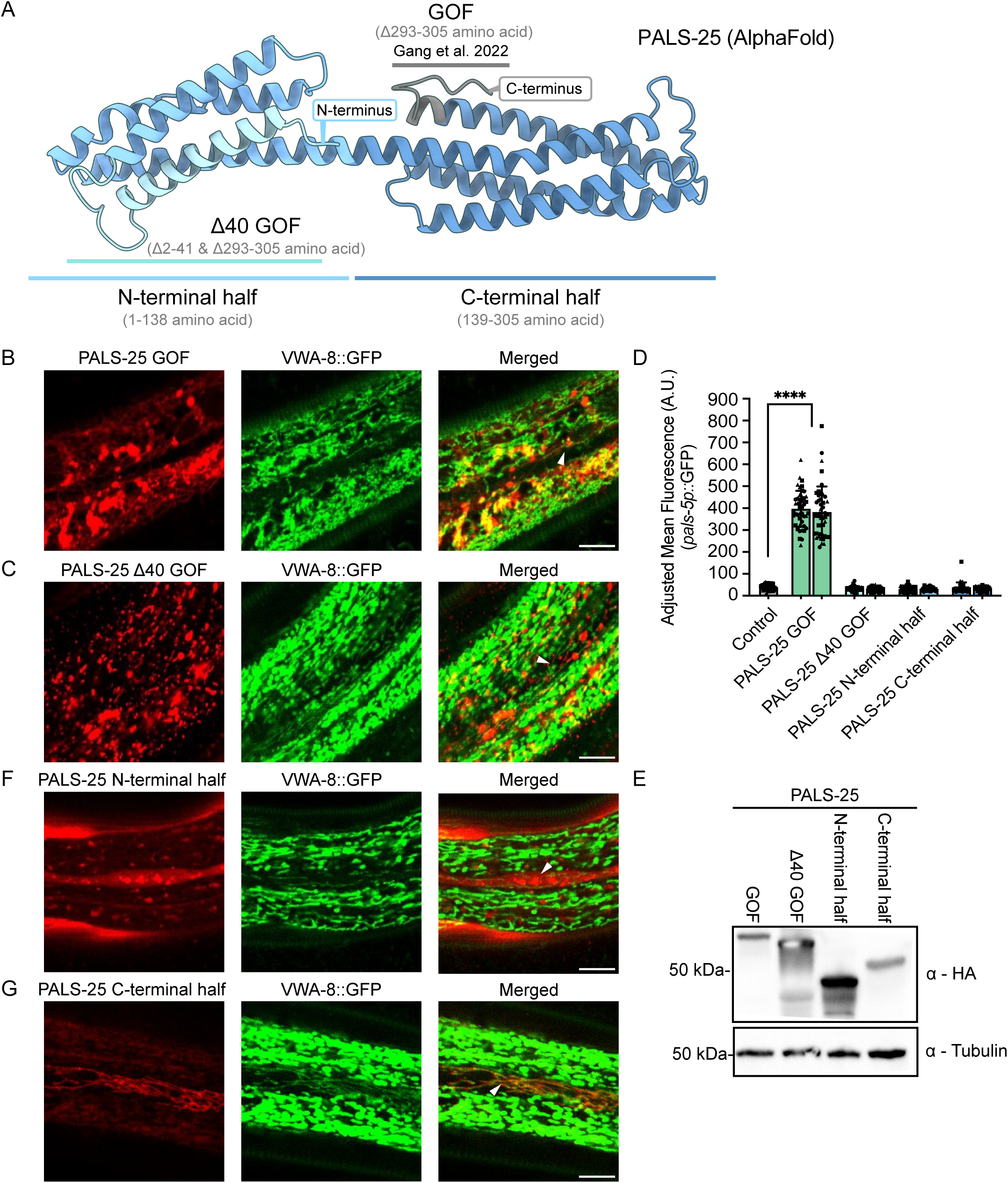
The C-terminal half of PALS-25 is required for mitochondrial localization, and N-terminal residues are required for signaling. **A)** Predicted structure of PALS-25 (AlphaFold DB Q3V5H4). Gray corresponds to the PALS-25 C-terminal 13 amino acids, which are deleted in PALS-25(GOF), causing PALS-22 dissociation and IPR induction (Gang et al. 2022). **B)** mScarlet::PALS-25 GOF localization with the mitochondrially localized protein VWA-8::GFP. **C)** mScarlet::PALS-25 τι40 GOF localization with the mitochondrially localized protein VWA-8::GFP. **D)** Quantification of *pals-5p::*GFP expression normalized to body area in adults (72 h post-L1). *pals-22 pals-25(jy80); jyIs8* animals (control) show low fluorescence and two positive control lines expressing mScarlet::PALS-25 GOF show significant induction of IPR reporter. n = 3 independent experimental replicates with 15 animals per genotype per experiment. *\*\*\*\* p <* 0.0001, one-way ANOVA with Dunnett’s multiple comparisons test. The graph shows mean values, with error bars represent standard deviations. Each symbol represents an individual animal. Different symbol shapes represent animals imaged on different days. **E)** Western blot analysis of different transgenically expressed PALS-25 mutant proteins in total protein lysates using commercially available HA antibody and tubulin antibody as a loading control. **F)** mScarlet::PALS-25 N-terminal half has diffuse cytosolic expression. **G)** mScarlet::PALS-25 C-terminal half has mitochondrial localization. For **B, C, F,** and **G,** Scale bar = 10 μm. White triangles denote seam cells. All strains tested in Figure 3 are in a *pals-22 pals-25(jy80)* mutant background.

Because many proteins are directed to mitochondria via a mitochondrial leader sequence (MLS) in their N-termini, we examined the N-terminus of PALS-25 for potential MLS sequences. There was not a significant hit for an MLS using prediction programs MitoFates^21^ and DeepMito^21^. To experimentally examine the possibility of an MLS, we deleted the N-terminal 40 amino acids of PALS-25, and then investigated subcellular localization. Here we saw that mScarlet::PALS-25(τι40 GOF) had a localization pattern similar to the PALS-25(GOF) (Fig 3B, C). Thus, the N-terminal 40 amino acids of PALS-25 do not appear to be responsible for mitochondrial localization. However, we found the N-terminal 40 amino acids are required for IPR induction. In contrast to the strong induction of the IPR reporter *pals-5p::*GFP in mScarlet::PALS-25(GOF) animals, mScarlet::PALS-25(τι40 GOF) had no *pals-5p::*GFP induction (Fig 3D). We confirmed that mScarlet::PALS-25(GOF) and mScarlet::PALS-25(τι40 GOF) proteins are expressed at similar levels, and thus lower PALS-25 expression levels are not responsible for the lack of IPR induction upon loss of the N-terminal 40 amino acids (Fig 3E).

To more broadly examine which region of PALS-25 directs localization to the mitochondria, we took advantage of AlphaFold, which predicts that PALS-25 protein structure is composed of an N-terminal bundle of 4 alpha-helices, and a C-terminal bundle of 4 alpha-helices (Fig 3A). We thus generated tagged versions of the N-terminal half and C-terminal half of PALS-25. Specifically, we fused mScarlet to the C-terminus of the N-terminal half of PALS-25 (1-138 aa), and we fused mScarlet to the N-terminus of the C-terminal half of PALS-25 (139-305 aa). Here we found that the N-terminal half of PALS-25 had a diffuse cytosolic localization pattern that did not colocalize with the mitochondrial marker VWA-8::GFP ^22^ (Fig 3F), whereas the C-terminal half of PALS-25 colocalized well with VWA-8::GFP (Fig 3G). Of note, neither the N-terminal or C-terminal half of PALS-25 could activate the IPR independently (Fig 3D). Altogether, these results indicate that the C-terminus of PALS-25 is both necessary and sufficient for localization to mitochondria. Furthermore, the N-terminal 40 amino acids are required for IPR induction.

### Loss of *pals-22* causes mitochondrial fragmentation

Upon analysis of *pals-22* RNAi-treated animals, we noticed a change in the morphology of mitochondria (Fig 2B, Supp Fig 4A). We quantified these effects in two *pals-22* loss-of-function mutants, *pals-22(jy1)* and *pals-22(jy3)* (Supp Fig 1C), which have similar levels of constitutive IPR induction, pathogen resistance, increased thermotolerance, as well as other phenotypes^11,16^. We first analyzed epidermal mitochondria using *col-19p::mito-GFP* (Fig 4A-C). Here, we found that *pals-22* mutants had a significantly lower mitochondrial form factor (Fig 4D), which is a metric for the complexity of mitochondria (see Methods). A higher form factor indicates more elongated mitochondria, and a lower form factor indicating more spherical mitochondria. In particular, epidermal mitochondria in *pals-22* mutants had significantly fewer branches and shorter branches, as well as slightly wider branch diameter (Fig 4E-G).

**Figure 4.**
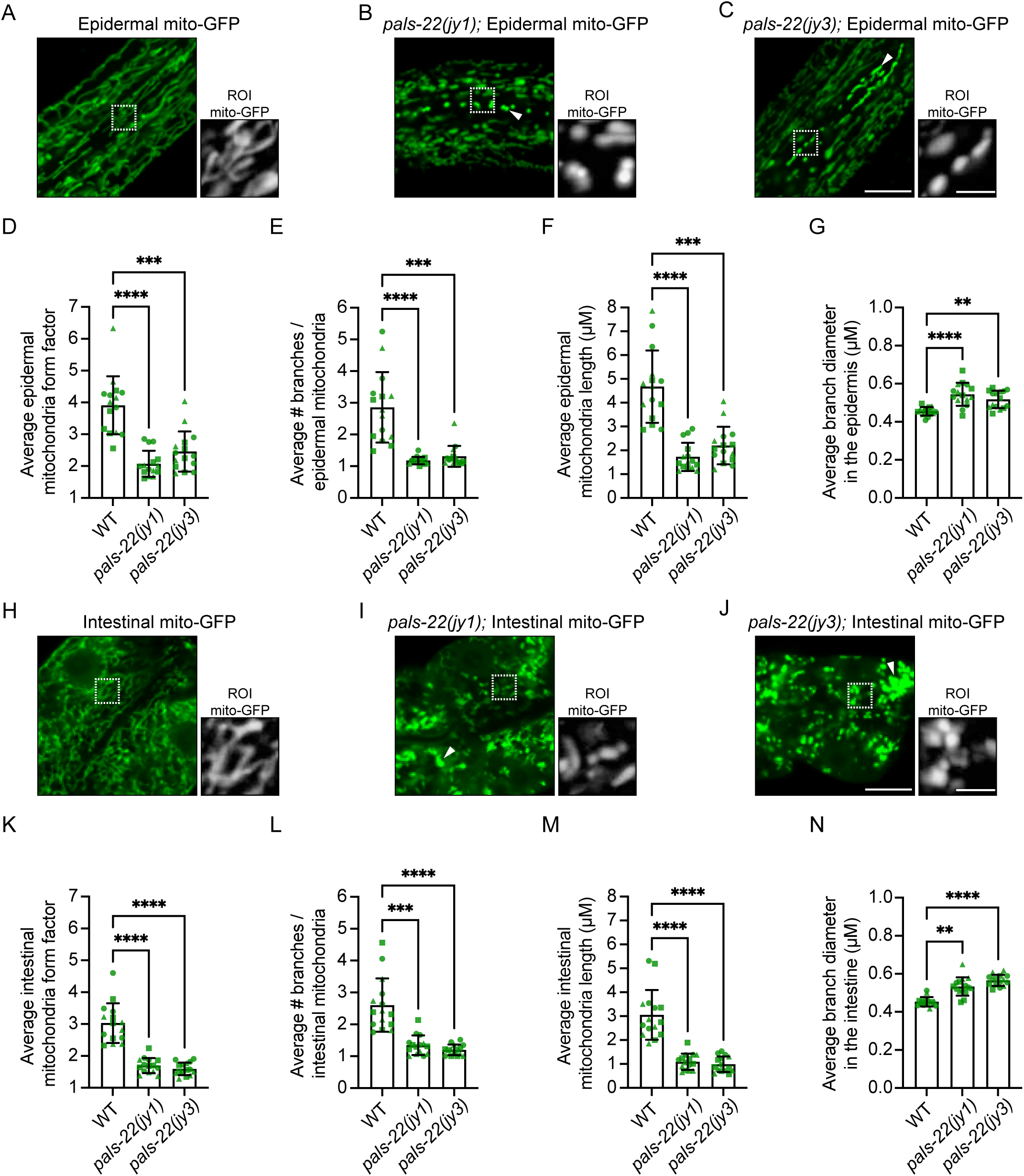
*pals-22* mutants have mitochondrial fragmentation. **A-C)** Representative images of wild-type **(A)**, *pals-22(jy1)* **(B)**, and *pals-22(jy3)* **(C)** young adult animals expressing *jyEx4796[col-19p::*mito-GFP] in the epidermis. **D-G)** *pals-22* mutants display altered mitochondria morphology in the epidermis, including decreased form factor (less elongated and more spherical morphology) **(D)**, fewer branches per mitochondria **(E)**, decreased average length **(F)**, and increased average diameter **(G)**. **H-J)** Representative images of wild-type **(H)**, *pals-22(jy1)* **(I)**, and *pals-22(jy3)* **(J)** young adult animals expressing *mgIs48[ges-1p::*mito-GFP] in the intestine. Scale bar = 10 μm. **K-N)** *pals-22* mutants display altered mitochondria morphology in the anterior intestine, including decreased form factor **(K)**, fewer branches per mitochondria **(L)**, decreased average length **(M)**, and increased average diameter **(N)**. For **A-C** and **H-J**, Scale bar = 10 μm. The white boxes indicate a region of interest (ROI) of the epidermal seam cells, and increased magnification of mitochondria morphology in the ROI based on mito-GFP expression is shown to the right. ROI scale bar = 2.5 μm. The exposure time was set to optimize mito-GFP expression in wild-type animals. However, we noted many *pals-22* mutants that displayed increased mito-GFP fluorescence intensity (white triangles). For **D-G** and **K-N**, ** *p <* 0.01, *** *p <* 0.001, *\*\*\*\* p <* 0.0001, Kruskal-Wallis test with Dunn’s multiple comparisons test. n = 15 ROIs analyzed per genotype, one ROI per animal, across three independent experimental replicates. Graphs show mean values, and error bars represent standard deviations. Different symbol shapes represent animals imaged on different days.

We next performed analysis of intestinal mitochondria as imaged by *ges-1*p::mito-GFP, and saw similar effects in this tissue; i.e. mitochondria in *pals-22* mutants had a lower form factor, fewer branches and shorter branches, and slightly wider branch diameter (Fig 4H-N). As expected, we observed these morphology changes were reversed to a wild-type phenotype in *pals-22/25(jy80)* double mutants (Supp Fig 4B), indicating that the effects on mitochondrial morphology of *pals-22* loss were dependent on *pals-25*. Therefore, loss of *pals-22* leads to broad changes in mitochondrial morphology in both the intestine and the epidermis, with a switch to mitochondria becoming more spherical, less branched, and less elongated, collectively described as being more fragmented.

To investigate how *pals-22*-mediated mitochondrial fragmentation might connect with known mitochondrial regulatory factors, we compared genes induced as part of the IPR to genes induced by the transcription factor ATFS-1, which mediates induction of the mitochondrial unfolded protein response (mitoUPR)^23–25^. Here we found some overlap between these gene sets, but the majority of mitoUPR genes are not part of the IPR, including canonical mitoUPR genes such as *hsp-6* and *hsp-60* (Supp Fig 4C). We next investigated whether knock-down of the mitochondrial fission factor *drp-1* might restore normal morphology in *pals-22* mutants. Of note, while loss of *drp-1* causes the outer mitochondrial membrane to be more elongated (Fig 1C), it causes fragmentation of the mitochondrial matrices (Supp Fig 4D)^19^. In *pals-*22 mutants, we found not only a lack of rescue by *drp-1* RNAi, but instead an exacerbation of the mitochondrial morphology phenotype, with extremely compacted mitochondria when *drp-1* RNAi knock-down was performed (Supp Fig 4E). In addition, we did not see rescue of the *pals-22* phenotype with RNAi against *pink-1*, which is required for mitophagy (Supp Fig E)^18^. Thus, the fragmentation of mitochondria in *pals-22* mutants appears to be distinct from changes caused by several canonical mitochondrial regulatory factors.

### IPR activation precedes mitochondrial fragmentation, which promotes defense against microsporidia infection

We next investigated whether PALS-22 was exerting its effects on mitochondrial morphology directly due to its localization to that organelle, or indirectly through the transcriptional IPR program. To distinguish between these models, we performed kinetic analysis to determine whether loss of PALS-22 leads to mitochondrial morphology changes before or after IPR induction. We used a PALS-22::AID strain, which has a degron tag that enables controlled degradation of this protein upon exposure to auxin^15,26^. With a time-course of auxin treatment, we found that after 3 hours of auxin treatment, there was robust induction of the IPR reporter *pals-5p::*GFP (Fig 5A, Supp Fig 5), whereas significant mitochondrial fragmentation in the intestine did not occur until 4 hours and later (Fig 5B,C). Similarly, *pals-5p::*GFP induction preceded mitochondrial fragmentation in the epidermis as well (Fig 5D-G). Specifically, there was no significant effect on mitochondrial form factor at 3 hours, but there was at 48 hours.

**Figure 5.**
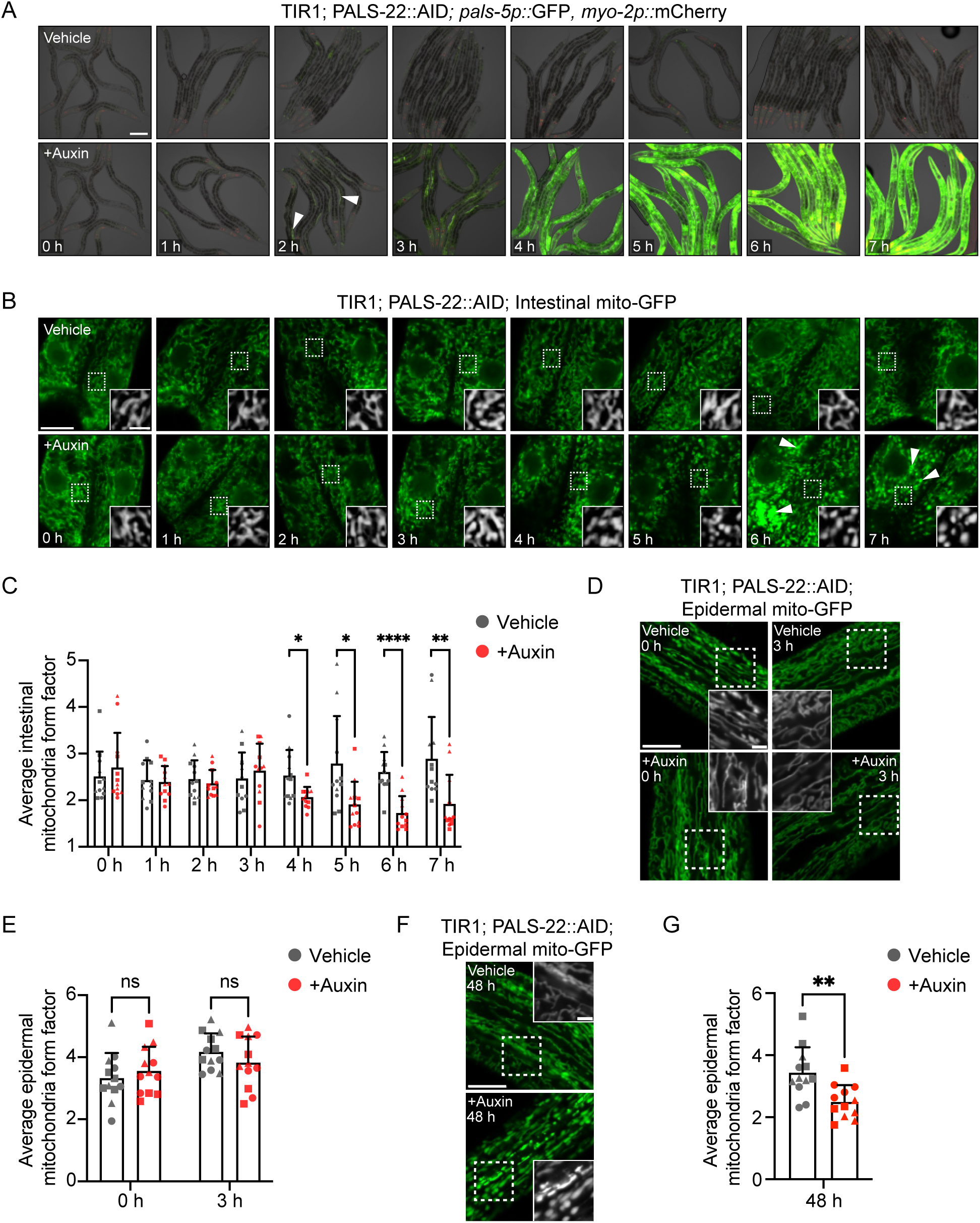
IPR reporter induction precedes mitochondrial fragmentation upon depletion of PALS-22. **A)** Time course images of transgenic animals expressing ubiquitous TIR1, endogenous *pals-22* tagged with AID, and the *pals-5p::*GFP IPR reporter treated with either ethanol vehicle control (top) or auxin (bottom) starting at 40 h post-L1. Auxin-mediated depletion of PALS-22 induces *pals-5p*::GFP IPR reporter expression as early as ∼2 h after auxin treatment (white triangles) and in most tissues by 3 h. *myo-2p::*mCherry is a marker for the transgene expressed in the pharynx. Images are an overlay of DIC, GFP, and mCherry fluorescence channels. Scale bar = 100 μm. **B)** Time course images of transgenic animals expressing ubiquitous TIR1, endogenous *pals-22* tagged with AID, and *mgIs48[ges-1p::mito-GFP]* treated with either vehicle control (top) or auxin (bottom) starting at 40 h post-L1. The white boxes indicate an ROI in the first ring of intestinal cells, and increased magnification showing mitochondria morphology in the ROI based on mito-GFP expression is inset. The exposure time was set to optimize mito-GFP expression at the 0 h treatment time point. However, we noted that increased time exposed to auxin caused increased mito-GFP fluorescence intensity (white triangles). **C.** Quantification of the treatment time course in **B.** Auxin-mediated depletion of PALS-22 induces significant changes in intestinal mitochondria morphology, as measured by form factor, starting 4 h after auxin treatment. **D-G)** Mitochondrial morphology analysis of *jyEx4796[col-19p::mito-GFP]* in the epidermis following auxin-mediated depletion of PALS-22**. D** and **F)** Representative images of 0 h and 3 h **(D)** or 48 h **(F)** treatment timepoints of mitochondria in the epidermis positioned relative to anterior intestinal cells (located above or below the first two rings of intestinal cells). The white boxes indicate an ROI, and increased magnification showing mitochondria morphology in the ROI based on mito-GFP expression is inset. **E)** Quantification of mitochondria form factor in the epidermis at 0 h vs 3 h treatment in **D**. Auxin-mediated depletion of PALS-22 does not induce changes in epidermal mitochondria morphology after 3 h. **G)** Quantification of mitochondria form factor in the epidermis at 48 h treatment in **F**. Auxin-mediated depletion of PALS-22 induces significant changes in epidermal mitochondria morphology 48 h after auxin treatment. For **B, D,** and **F**, scale bar = 10 μm, and ROI scale bar = 2.5 μm. For **C, E,** and **G**, ns = not significant, *\* p <* 0.05, *\*\* p <* 0.01, *\*\*\*\* p <* 0.0001, Welch’s t-test for vehicle control vs. auxin at each time point. n = 12 ROIs analyzed per time point and condition, two ROIs per animal, across three experimental replicates. Graphs show mean values and error bars represent standard deviations. Different symbol shapes represent ROIs from animals imaged on different days.

To test the model that loss of *pals-22* causes mitochondrial fragmentation indirectly through IPR induction, we activated the IPR in a distinct manner. Specifically, we performed RNAi against *pals-17*, which encodes a negative IPR regulator that localizes to the plasma membrane in the intestine, without obvious localization to mitochondria^12^. Here, we found that RNAi against *pals-17* had similar effects to the loss of *pals-22,* with intestinal mitochondria becoming fragmented (Supp Fig 6). This finding, together with the PALS-22 depletion studies mentioned above, supports the model that the *pals-22/25* module controls mitochondrial fragmentation indirectly through IPR induction.

We next asked whether *N. parisii* associates with *C. elegans* mitochondria and found, via transmission electron microscopy (TEM), that it does (Fig 6A). This type of association has been seen with microsporidia species infecting other hosts^27^, and is thought to aid in ATP and other nucleotide transport into microsporidia, which lack their own mitochondria and nucleotide biosynthesis pathways. To determine whether host mitochondrial fragmentation may regulate resistance against microsporidia, we induced mitochondrial fragmentation through either *drp-1* RNAi or *fzo-1* RNAi and then measured *N. parisii* pathogen load. In both cases we found that *C. elegans* had increased resistance to infection (Fig 6B). Notably, neither of these RNAi treatments caused induction of the *pals-5*p::GFP reporter (0% of animals had induction, n > 200 animals examined for each condition). Thus, fragmentation of the mitochondrial matrix independent of *pals-5*p::GFP induction appears to protect *C. elegans* against microsporidia infection.

**Figure 6.**
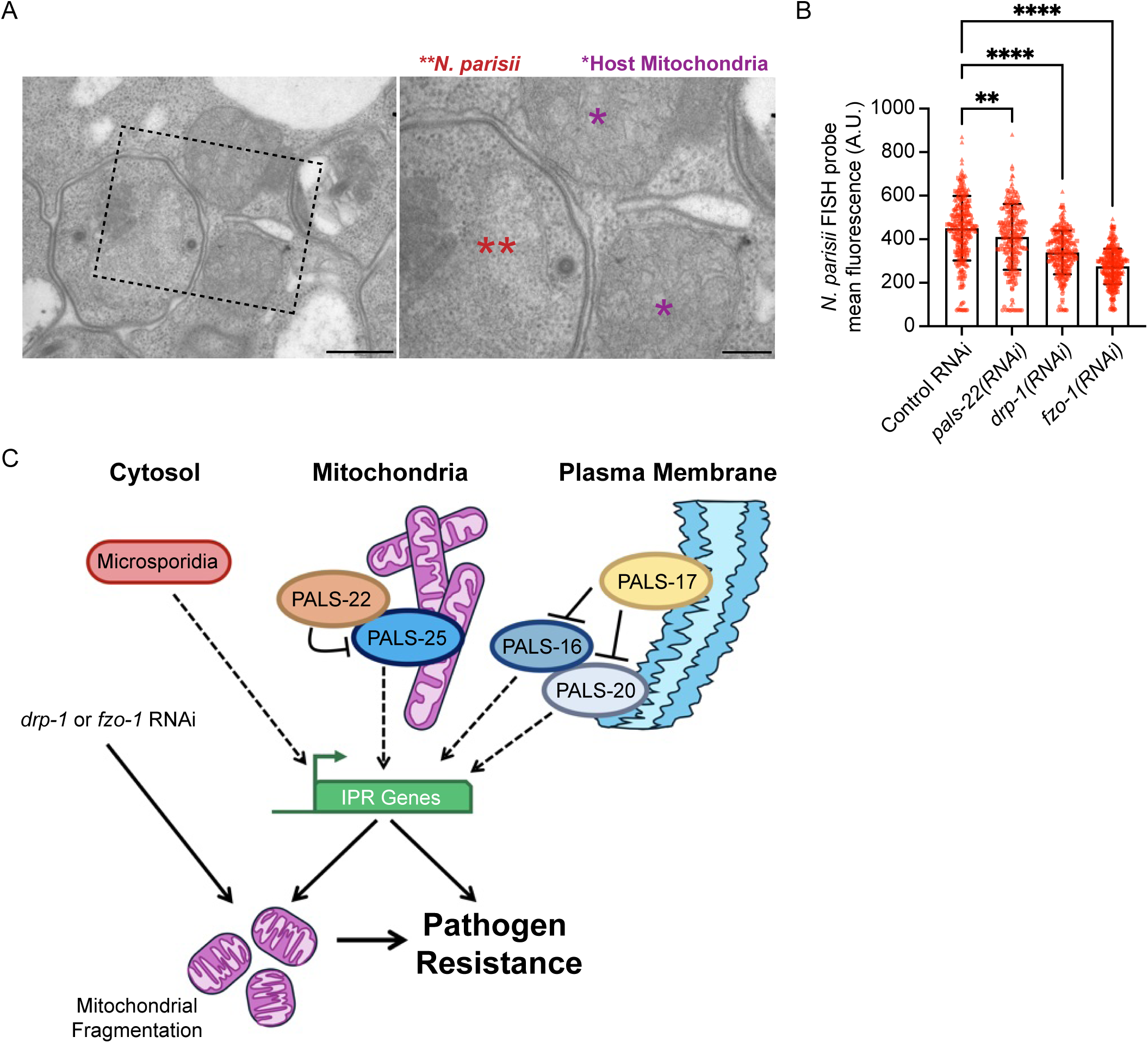
Animals with mitochondrial fragmentation independent of IPR induction have decreased *N. parisii* pathogen load. **A)** Representative TEM images of *N. parisii* sporonts (red double asterisks) associated with *C. elegans* mitochondria (purple asterisks) during infection. Scale bar = 500 nm, ROI scale bar = 200 nm. **B)** Wild-type animals treated for two generations on *drp-1* or *fzo-1* RNAi display increased resistance to *N. parisii* infection. ** *p* < 0.01, **** *p* < 0.0001, one-way ANOVA with Dunnett’s multiple comparisons test. n = 209-279 animals per treatment across three experimental replicates. The graph shows mean values, and error bars represent standard deviations. Different symbol shapes represent animals analyzed in distinct infection experiments. **C)** Model for PALS-22 and PALS-25 as regulators of mitochondrial fragmentation through IPR activation. Mitochondrial fragmentation, either through IPR activation or other morphology regulators, promotes resistance to microsporidia infection.

## Discussion

The immune switch proteins PALS-22 and PALS-25 cause a dramatic rewiring of *C. elegans* physiology, controlling the balance between an immune state and a growth state^11^. However, PALS proteins lack clear functional domains and have a poorly understood mechanism of action. Here we show that both PALS-22 and PALS-25 proteins localize to mitochondria, apparently at the outer membrane, with PALS-22 being dependent on PALS-25 for localization to mitochondria. We define the C-terminal half of PALS-25 to be necessary and sufficient for mitochondrial localization, and the N-terminal 40 amino acids as being required to activate the IPR. PALS-22-mediated control of mitochondrial fragmentation occurs after activating the IPR transcriptional program, and can be induced by loss of another IPR inhibitor PALS-17, suggesting that IPR activation may trigger fragmentation (Fig 6C).

Two recent studies in mammalian tissue culture cells have shown that microsporidia infection promotes fragmentation of host mitochondria^28,29^. During infection, it can be difficult to disentangle the role of host vs pathogen-derived factors in driving fragmentation. One of these studies suggested that microsporidia infection activated DRP-1, which caused mitochondria fragmentation to ultimately aid microsporidia growth, based on analysis with the mdivi-1 inhibitor, which has multiple impacts in the cell^28,30^. In contrast, here we report that mitochondrial fragmentation appears to impair microsporidia growth (Fig 6C). Further studies will be needed to determine when and where mitochondrial fragmentation is beneficial vs inhibitory for microsporidia growth.

Several external stressors cause fragmentation of mitochondria, such as wounding of the *C. elegans* epidermis, where fragmentation appears to promote wound healing^31^. Similarly, heat shock can induce fragmentation of mitochondria, which in some paradigms appears to then promote resistance to heat shock^32,33^. Fragmentation is required for subsequent removal of damaged mitochondria through mitophagy. Similar themes have been observed after *C. elegans* infection with the extracellular bacterial pathogen *Pseudomonas aeruginosa*, where mitochondrial mass is reduced^34^.

In contrast to stressors like heat shock and infection that trigger fragmentation of mitochondria, upregulation of many stress/immune response pathways trigger hyperfusion or expansion of mitochondrial networks. For example in *C. elegans*, ATFS-1, the transcription factor that regulates the mitoUPR, appears to promote expansion of mitochondrial networks^35^, and overexpression of heat shock factor 1 (HSF1) leads to hyperfusion of mitochondria and increased lifespan^36^. Wild-type function of the bZIP transcription factor ZIP-2, which promotes defense against *P. aeruginosa*, also promotes mitochondrial fusion^37^. Similarly, in mammals, activation of the endoplasmic reticulum stress pathway causes mitochondrial hyperfusion^38^, and activation of the RIG-I-like receptor pathway can also activate hyperfusion, through separate kinase driven mechanisms^39^.

Our observation that the IPR triggers fragmentation in the absence of a stressor is perhaps unusual for a stress response pathway, as the pathways mentioned above promote mitochondrial fusion. One exception is the transcription factor HLH-30/TFEB, which promotes mitochondrial fusion in a cell-autonomous manner, but mitochondrial fragmentation in a cell non-autonomous manner^40^. Similarly, mitochondrial fragmentation in neurons was found to cell non-autonomously cause fragmentation in the intestine ^41^. Of note, fragmented mitochondria may switch to fatty acid oxidation to provide fuel, which perhaps is beneficial in the context of intracellular pathogens like microsporidia^42^. Indeed, there are indications that microsporidia cause a decrease in host fat levels and rely on host fatty acids, and thus depleting fatty acids might aid in resistance to infection^43,44^.

We found that loss of either PALS-22 or PALS-17 would trigger mitochondrial fragmentation. In contrast to PALS-22 and PALS-25, the protein PALS-17 and its positive regulator PALS-20, which another OFF/ON switch of the IPR, localize to the apical membrane of intestinal cells^12^. It is possible that membrane localization is critical for mechanism of action for these PALS proteins.

Many other immune regulators are localized to membranes, such as the viral RNA sensor RIG-I, which localizes to mitochondria upon activation and binding to its downstream signaling partner MAVS. More detailed analysis of the activity of PALS-22/25 and other PALS proteins at mitochondrial and other membranes will help decipher how they trigger downstream IPR signaling and immunity.

## Materials and Methods

### *C. elegans* growth and maintenance

*C. elegans* worms were maintained following standard methods^45^ at 20 °C on Nematode Growth Media (NGM) plates seeded with streptomycin-resistant *Escherichia coli* OP50-1 unless stated otherwise for specific experiments. The strains used in this study are described in Supp Table 1.

### Synchronization of *C. elegans* growth

To obtain populations of worms synchronized at the same life stage, gravid adults were washed from NGM + OP50-1 plates with M9 media into a 15 ml conical tube. The contents of the tube were then centrifuged at 3,000 rpm for 30 s to pellet the worms, and the supernatant was removed, leaving ∼2 ml of M9 with the worm pellet. 800 µl of 5.65-6% sodium hypochlorite solution and 200 µl of 2 M NaOH were added, and the tube was vortexed intermittently for ∼2 minutes or until the majority of adult animals had partially dissolved and released embryos into solution. The embryos were washed by filling the tube to 15 ml with M9, the tube was centrifuged to pellet the embryos, and the supernatant was removed. The 15 ml M9 wash-and-centrifuge process was repeated 4 additional times (5 washes total), and the embryos were resuspended to a final volume of ∼5 ml in M9. The tube was placed in a 20 °C incubator with continual rotation for 16-24 h to hatch synchronized L1s.

### RNA interference

RNAi was performed using the *E. coli* HT115 feeding method. L4440 control vector, *pals-22*, *pals-25*, *pals-17*, *fzo-1*, *eat-3*, *pink-1*, and *drp-1* RNAi clones in the L4440 vector background were used. Colonies for each clone were grown overnight in LB broth supplemented with 50 µg/ml carbenicillin at 37 °C with shaking at 250 rpm. 6-cm or 10-cm RNAi plates (NGM plates supplemented with 5 mM IPTG and 1 mM carbenicillin) were seeded with 400 µl or 1 ml of overnight culture, respectively, and were dried in a biosafety cabinet. The plates were then stored at room temperature in the dark for 48-72 h to allow the RNAi bacteria lawn to grow. The RNAi knockdown procedure for specific experiments is described below.

### Confocal Imaging

For all confocal imaging, all animals were mounted on a 5% agarose pad in 100 mM levamisole and imaged on an LSM700 confocal microscope with Zen 2010 software.

### Imaging PALS-22/25::GFP, Mitochondria, and Co-localization of PALS-22/25

For Fig. 1A,B and Fig. 2: Synchronized L1 animals expressing either *jyEx193[pals-22p::pals-22::GFP::3xFLAG]* or *jyEx237[pals-22p::pals-25::GFP::3xFLAG]* were plated to 6-cm RNAi plates seeded with either L4440 control vector, *pals-22(RNAi)*, or *pals-25(RNAi)* and were incubated at 20 °C for 72 h. MitoTracker Red CMXRos (Invitrogen M7512) staining of the *C. elegans* epidermis was performed using a 1 mM MitoTracker stock solution in DMSO diluted to 5 µM in M9, and 200 µl of 5 µM MitoTracker was then aliquoted into a 1.5 ml microcentrifuge tube for each strain and/or RNAi condition. ∼50 adult animals per strain/condition were picked into 5 µM MitoTracker and were incubated in the dark at room temperature for 10 min with mixing by intermittent inversion. The worms were then pelleted by gravity settling, the supernatant was removed, 1 ml of M9 was added to the tube, and the worms were gently vortexed. The gravity settling and M9 wash procedure was repeated twice (3 washes total), and the worms were transferred to 10-cm NGM + OP50-1 plates. The worms were left to recover from MitoTracker treatment for 2 h in the dark at 20 °C. Localization of PALS-22::GFP or PALS-25::GFP with MitoTracker Red in the epidermis following the different RNAi treatments was assessed with confocal imaging. For Fig. 1C: Synchronized L1 animals expressing either *jyEx193[pals-22p::pals-22::GFP::3xFLAG]* or *jyEx237[pals-22p::pals-25::GFP::3xFLAG]* were plated to 6-cm RNAi plates seeded with either L4440 control vector, *fzo-1(RNAi)*, *eat-3(RNAi)*, or *drp-1(RNAi)* and were incubated at 20 °C for 72 h. Localization of PALS-22::GFP or PALS-25::GFP in the epidermis following the different RNAi treatments was assessed by confocal imaging. For Fig. 1D: L4 animals were picked from a mixed stage population of ERT1255 grown on 6-cm NGM + OP50-1 plates and co-localization was assessed by confocal imaging.

### Molecular Cloning and transgenesis

Four plasmids were generated for use in this study: pET787, pET789, PET792, and pET793. pET787[*pals-22p::gfp::pals-22::3xflag::sl2Intergenic::3xHA::wrmScarlet::pals-25::unc-54-3’utr*] was generated using Gibson assembly of gBlocks containing *gfp::pals-22::3xflag::sl2Intergenic* and *sl2Intergenic::3xHA::wrmScarlet::pals-25* with a vector backbone containing *pals-22p* and *unc-54* 3’UTR. pET789 was generated through a Q5 deletion protocol using pET729 as a template^15^, deleting residues 2-41. pET792 was generated by Q5 deletion protocol using pET790 [*pals-22promoter::pals-22cDNA::sl2::pals-25cDNA::wrmScarlet::3xHA::unc-54 3’UTR*] deleting residues 139-305. pET793 was generated by deletion of pET791[*pals-22p::pals-22::3xflag::sl2Intergenic::3xHA::wrmScarlet::pals-25::unc-54-3’utr*] deleting residues 2-138. All plasmids were sequence confirmed using Primordium. Gibson assembly protocol volume calculator was obtained at (https://barricklab.org/twiki/bin/view/Lab/ProtocolsGibsonCloning). DNA for injection by was prepared by overnight culture of 4 ml LB + 100 ng/µL carbenicillin and cultures were grown for ∼17 hours. This overnight culture was miniprepped using the Qiagen QIAprep Spin Miniprep Kit following manufacturer protocols with the exception that no RNAse was added to buffer P1. Injection mixes were prepared to a final total [DNA] of 100 ng/µl in 10 µl. 0.5 µl of the mix was loaded into microinjection needles and injected into gravid adult gonads. pET787 was injected at a concentration of 20 ng/µl and co-injected with a green neuronal marker (*gcy-8p::GFP*) at 20 ng/µl and pBluescript was added to a final concentration of 60 ng/µl. pET789 was injected at a concentration of 60 ng/µl and co-injected with a green neuronal marker (*gcy-8p::GFP*) at 20 ng/µl and pBluescript was added to a final concentration of 20 ng/µl. pET792 and pET793 were injected at a concentration of 20 ng/µl and co-injected with a green neuronal marker (*gcy-8p::GFP*) at 20 ng/µl and pBluescript was added to a final concentration of 60 ng/µl. Transgenic lines were identified green neuronal marker. For Fig 3C-G: At least two different F1 transgenic lines were isolated per injection mix. For Fig 3B: pET729 was injected at a concentration of 60 ng/µl and co-injected with a green neuronal marker (*gcy-8p::GFP*) at 20 ng/µl, and pBluescript was added to a final concentration of 20 ng/µl.

### Imaging Colocalization of PALS-25 fragments with mitochondria

Approximately 1-day old adult animals from a mixed stage population of ERT1368, 1369, 1372, 1373 grown on 6-cm NGM + OP50-1 plates were imaged, with co-localization assessed by confocal imaging on a Zeiss LSM700.

### RNA extraction and RT-qPCR

For Fig. S2A,B: ∼3000-3,500 synchronized L1s for ERT964 *pals-22 pals-25(jy80); jyEx193[pals-22p::pals-22::GFP::3xFLAG]* or ERT965 *pals-22 pals-25(jy80); jyEx237[pals-22p::pals-25::GFP::3xFLAG]* were plated to 10-cm RNAi plates seeded with 1 ml overnight culture of L4440 control vector or *pals-22* RNAi clones and were incubated at 20 °C for 48 h. For Fig. S2C: ∼1000 synchronized L1s of ERT714 *pals-22 pals-25(jy80)*, ERT751 *pals-25(jy111)*, ERT964 *pals-22 pals-25(jy80) unc-119(ed3) III; jyEx193[PALS-22::EGFP::3xFLAG, unc-119(+)]*, and ERT965 *pals-22 pals-25(jy80) unc-119(ed3) III*; *jyEx237[PALS-25::EGFP::3xFLAG, unc-119(+)]* were plated onto 2 × 10-cm plates and incubated at 20 °C for 50 h. For all RT-qPCR experiments, RNA was extracted with TRI reagent (Molecular Research Center TR118), isolated with BCP reagent (Molecular Research Center BP151), washed with 100% isopropanol followed by 75% ethanol, and resuspended in HyClone nuclease-free H_2_O (Cytiva SH30538.03). cDNA was synthesized from total RNA using iScript (Bio-Rad 1708890). qPCR was performed with iQ SYBR Green Supermix (Bio-Rad 1708880) with a CFX Connect Real-Time PCR Detection System (Bio-Rad). Expression values for genes of interest were normalized to the expression of a control gene, *snb-1*, which does not show altered expression following IPR activation^46^. The Pffafl method was used for quantification^47^. Primers used for RT-qPCR analysis are provided in Supp Table 1.

### Protein collection

For Fig. S3B,C: ∼1500-2000 synchronized L1s were plated onto RNAi plates seeded with either L4440 control vector, *pals-22*, or *pals-25* RNAi and were incubated at 20 °C for 72 h. Worms were washed twice with ice-cold PBS + 0.1% Tween-20 (PBS-T), and following the final wash, gravity settled on ice. A 15 µl aliquot of the dense worm pellet was then transferred to a 1.5 ml microcentrifuge tube and was combined with 20 µl of 4x Laemmli sample buffer (Bio-Rad #1610747) supplemented with 200 mM dithiothreitol (DTT), 25 µl ice-cold PBS-T, and mixed by vortexing. The tubes were incubated at 100 °C for 15 minutes to digest the worms with gentle vortexing at each 5-minute interval. The protein lysates were frozen at -30 °C prior to SDS-PAGE and Western blot. For Fig. 3E: PALS-25 fragment expressing strains, 2 × 10-cm plates containing 15 transgenic positive L4s were plated. 72 hours after plating P0, 1 ml of 10x concentrated OP50 was added to each plate. 24 hours later, worms were washed off the plates with M9 + 0.1% Tween-20 (M9T) into a 15 ml conical tube, centrifuged at 3,000 rpm for 30 s to pellet the worms, and the supernatant was removed. Animals were washed twice with 15 ml of M9T, centrifuging at 3,000 RPM for 30 s to pellet worms and remove supernatant each time. After the final wash, supernatant was aspirated down to 1 ml and worms + M9T was transferred to a microcentrifuge tube and worms were then spun in a minifuge for 20 seconds. Supernatant was aspirated down to 100 µl. 33 µl of 4x Laemmli sample buffer (Bio-Rad #1610747) supplemented with 400 mM dithiothreitol (DTT) was mixed by vortexing. The tubes were incubated at 100 °C for 15 minutes to digest the worms with gentle vortexing at each 5-minute interval. The protein lysates were frozen at -30 °C prior to SDS-PAGE and Western blot.

### Western blot analysis

For Fig. S3B-C: Protein lysates were thawed from -30 °C on ice, centrifuged at 13,000 rpm for 10 minutes, and 10 µl per sample were separated on a 4-15% sodium dodecyl sulfate– polyacrylamide gel electrophoresis (SDS-PAGE) precast gel (Bio-Rad #4561086) at room temperature. Proteins were transferred onto a polyvinylidene difluoride (PVDF) membrane at 4°C, 100 V, for 1.5 h. Blocking was performed in 5% nonfat dry milk dissolved in PBS-T for 2 h at room temperature. The PVDF membrane was then incubated with primary antibodies diluted in 5 ml blocking buffer overnight at 4 °C with rocking. Custom polyclonal anti-PALS-22 and anti-PALS-25 were developed and used at 1:1,000. Commercial monoclonal anti-α-tubulin produced in mice (Millipore Sigma, Cat#T9026) was used at 1:4,000. Following incubation in primary antibody, the PVDF membrane was washed three times in PBS-T, and then incubated with secondary antibodies conjugated with horseradish peroxidase in 5 ml blocking buffer for 1.25 h at room temperature with rocking (for anti-PALS-22 and anti-PALS-25 primary: goat anti-rabbit 1:10,000 (Millipore Sigma, Cat#401315-2ML) For anti-tubulin primary: goat anti-mouse 1:10,000 (Millipore Sigma, Cat#401215-2mL). The PVDF membrane was washed three times in PBS-T, then was treated with enhanced chemiluminescence (ECL) reagent (Cytiva RPN2209) for 5 minutes and imaged using a Chemidoc XRS+ with Image Lab software (Bio-Rad). Quantification of PALS-22::GFP or PALS-25::GFP band intensities was determined using Image Lab software (Bio-Rad), and each sample was normalized to its tubulin expression levels. For Fig. 3E: Protein lysates were thawed from -30 °C on ice, centrifuged at 15,000 rpm for 15 minutes, and 20 µl per sample were separated on an Any kD Mini-PROTEAN TGX Precast Protein Gel 10-wells with 7 µL of Precision Plus Protein Dual Color Standards reference band ladder. Protein standards and each sample was loaded twice. The gel was run at 120 V for 1 hour and 30 minutes. Proteins were transferred onto a polyvinylidene difluoride (PVDF) membrane at 4 °C, 30 V, for 18 h. The PVDF membrane was cut in half such that each half had a reference ladder and one lane for each sample. Blocking was performed as mentioned above. Each half of PVDF membrane was then incubated with primary antibodies diluted in 5 ml blocking buffer overnight at 4 °C with rocking (1:1000 for Anti-HA (Cell Signaling, Cat #C29F4), and 1:5000 for anti-tubulin). the PVDF membrane was washed three times in PBS-T and then incubated with secondary antibodies conjugated with horseradish peroxidase in 5 ml blocking buffer for 2 h at room temperature with rocking (for anti-HA: goat anti-rabbit 1:10,000; for anti-tubulin primary: goat anti-mouse 1:1,000). The PVDF membrane was washed and imaged as mentioned above.

### AlphaFold and ChimeraX visualization

AlphaFold DB version 2022-11-01, created with the AlphaFold Monomer v2.0 pipeline. ChimerX-1.4 was used to visualize .pdb files.

### Measuring IPR induction in PALS-25 fragment expressing strains

For Fig. 3D: ∼500 synchronized L1s were plated onto 2x plates for strains ERT930, ERT1008, ERT1010, ERT1331, ERT1332, ERT1334, ERT1335, ERT1301, ERT1303 and incubated at 20°C for 72 h. Plates were then washed off and pooled per genotype with 3 ml of M9T and collected into a 1.5 ml conical tube. Each tube was then washed 3 times with M9T, and allowed to gravity settle before aspirating down to 100 µl final volume. After final aspiration, 100 µl of M9T+ 10 uM levamisole was added. 96 clear bottom black well plates were used, and 4x wells per geneotype were filled with 200 µl of 10 uM levamisole, and animals were added to each well, optimizing volumes to obtain a high # of worms with little to no overlap of adult animals.

The final volume per well was 300 µl. Samples were analyzed for *pals-5p::*GFP expression on an ImageXpress Nano plate reader using the 4x objective (Molecular Devices, LLC, San Jose, CA) and MetaXpress 2018 software version 6.5.2.351. The worm area was traced using FIJI software (excluding the head regions due to *gcy-8p::GFP* co-injection marker expression), and the average fluorescence intensity of each worm was quantified with the background fluorescence of the well subtracted. 15 animals per genotype were quantified for each experimental replicate, and three independent experiments were performed.

### Imaging the mitochondria morphology of *pals-22* mutants

For Fig. 4: Wild-type animals expressing *mgIs48[ges-1p::mito-GFP]* or *juEx4796[col-19p::mito-GFP]* were grown for 48 h at 20 °C to the young adult life stage. *pals-22(jy1)* and *pals-22(jy3)* mutants, which are developmentally delayed^11^, expressing *mgIs48* or *juEx4796* were grown for 54 h at 20 °C to the young adult life stage. GFP-labeled mitochondria were assessed by confocal imaging. In the intestine, imaging was restricted to the first ring (four anterior-most cells) in the same focal plane as the intestinal nuclei determined by the absence of GFP signal within the nuclei. In the epidermis, images were collected by first identifying alae in the cuticle using transmitted light in the anterior half of the worm. Next, a focal plane underneath the alae was identified using GFP fluorescence such that seam cells and surrounding hyp7 cells had GFP-labeled mitochondria in focus. To minimize changes in mitochondrial network morphology as a result of prolonged immobilization on the agar pad and/or exposure to levamisole, images were only collected within 10 minutes after mounting before a new slide was prepared. Confocal imaging settings were independently optimized for each transgene based on the expression in wild-type controls. See below for morphology analysis. For Figure S4B: Wild-type, *pals-22(jy3)*, and *pals-22 pals-25(jy80)* adults were picked from 6-cm NGM + OP50-1 plates into a 1.5 ml microcentrifuge tube containing 200 µl of 2.5 µM MitoTracker Red diluted in M9. The worms were incubated in the dark for 10 min with mixing by intermittent inversion. The worms were then pelleted by gravity settling, the supernatant was removed, 1 ml of M9 was added to the tube, and the worms were gently vortexed. The gravity settling and M9 wash procedure was repeated twice (3 washes total), and the worms were transferred to 6-cm NGM + OP50-1 plates. The worms were left to recover from MitoTracker treatment for 2 h in the dark at 20 °C prior to confocal imaging.

### Imaging IPR activation and mitochondria morphology following auxin-mediated depletion of PALS-22

For Fig. 5 and Fig. S5: 6-cm NGM plates supplemented with either 200 µM auxin or vehicle control (0.15% ethanol) were seeded with 200 µl of 10x concentrated OP50-1 spread across the entire surface of the plate. For intestinal analysis, synchronized L1s of ERT1017 and ERT1187 were plated on NGM + OP50-1 and incubated at 20 °C for 40 h. A population of worms for each strain was then split and plated to auxin or vehicle treatments. Worms were then collected from each condition to collect a total of 8 time points (40-47 h post L1, 0-7 h on treatment). For epidermal analysis, synchronized L1s of ERT1224 and ERT1017 were plated to NGM + OP50-1 and incubated at 20 °C for 48 h or plated directly on auxin or vehicle plates. A population of worms for each strain plated on NGM + OP50-1 was split and plated on auxin or vehicle plates. All conditions were removed from their plates at specified time points: 0, 3, and 48 h on auxin. For all experiments, animals were removed immediately at specified time point and. were mounted and imaged for confocal imaging as described above. See below for mitochondrial morphology analysis.

### Mitochondrial morphology analysis in *pals-17* RNAi-treated animals

For Fig. S6: ∼250 synchronized L1s of ERT1145: *mgIs48[ges-1p::mito::GFP]* and ERT054: *jyIs8[pals-5p::gfp, myo-2p::mCherry] X* were each plated onto 4 × 6-cm OP50 plates and incubated at 20 °C for 20 h. For each strain, 2x plates were washed off and pooled with 2 ml of M9 and transferred into 1.5 ml conical tubes. The worms were then washed 3 times with M9, and pelleted down to 100 µl final volume. Each 1.5 ml conical was then split onto 2x plates of either *pals-17* RNAi or L4440 EV RNAi. Worms were then grown for 24 h. Worms were then imaged using the confocal as described above. At least five animals treated with *pals-17* RNAi and five animals treated with L4440 RNAi were imaged. Five animal samples were randomly selected per treatment, and two regions of interest were chosen from each animal. Downstream morphology analysis and quantification were performed using the protocol below.

### Mitochondria morphology analysis

Confocal images of intestinal and epidermal mitochondria labeled with GFP were analyzed with FIJI software^48^. For each animal, 10 µM × 10 µM square regions of interest (ROI) were selected for analysis. Several pre-processing steps in FIJI were performed to enhance mito-GFP signal to background noise, including, in order, a) subtract background, b) sigma filter plus, c) enhance local contrast, and d) gamma correction. The processed ROI was masked using the Mitochondria Analyzer plugin (https://github.com/AhsenChaudhry/Mitochondria-Analyzer/tree/master). Mitochondria Analyzer 2D analysis was performed on a per-ROI basis (i.e., morphology characteristics for each mitochondrion in the ROI were quantified and then averaged for the number of mitochondria detected in the ROI). For analysis of intestinal mitochondria, ROIs selected in the first ring were not cell-specific and, in some cases, spanned adjacent cells. Likewise, for analysis of epidermal mitochondria, ROIs spanned both seam cells and surrounding hyp7 in the anterior half of the animal. A step-by-step protocol for image pre-processing and analysis with Mitochondria Analyzer can be found at GitHub (https://github.com/TroemelLabUCSD/Mitochondria-and-PathogenLoad-Analysis), Fiji Macros linked here used for Fig 5C,E,G, S6C. The ‘Form Factor’ is a shape measurement where 1 equals a perfectly round 2D object and increases with object elongation.

### Imaging mitochondrial morphology in wild-type and *pals-22* mutants following RNAi mediated knockdown of mitochondrial morphology regulating factors

For Fig. S4A: ∼100 synchronized L1s of *juEx4796[col-19p::mito-GFP]* were plated on EV or *pals-22* RNAi plates seeded with 300 μl of RNAi culture. Mitochondria morphology was assessed with confocal imaging at 72 h; ; images were only collected within 10 minutes after mounting before a new slide was prepared. For Fig. S4D,E: ∼300 synchronized L1s of ERT1145 and ERT1146 were plated on EV, *drp-1*, or *pink-1* 6-cm RNAi plates seeded with 300 μl of RNAi culture, and mitochondrial morphology was assessed with confocal imaging at 48 and 72 h.

### Analysis of mitoUPR and IPR upregulated genes

Genes significantly upregulated (logFC > 0, p < 0.05) by the *atfs-1(et15)* gain-of-function allele, which constitutively activates the mitoUPR^24^, were compared to the 80 canonical IPR genes^11^ using WormBase IDs with the tool: https://bioinformatics.psb.ugent.be/webtools/Venn/.

### *N. parisii* transmission electron microscopy and pathogen load assays

*C. elegans* were infected with *N. parisii* spores at 25 °C and then fixed at the late meront/early sporont stage, approximately 40 hours post-inoculation for transmission electron microscopy as described^49^. To perform RNAi for Fig. 6B, overnight cultures were prepared at 30 °C for 19.5 h in 4 ml LB broth supplemented with µg/ml carbenicillin. RNAi plates were prepared for each experimental replicate by plating 750 μl of 4x concentrated RNAi overnight culture for L4440, *pals-22*, *drp-1,* and *fzo-1*. 15 L4 P0 N2 animals were plated onto 2x RNAi plates. 1 ml of 6x concentrated RNAi overnight culture that was grown at 30 °C for 19.5 h and induced with a final concentration of 2mM IPTG for 2 h at 30 °C was added to plates ∼72 h after plating P0s. F1 worms were then bleached/ synchronized ∼96 hours after plating P0s to obtain F2s. *N. parisii* spores were prepared as described previously^50^. ∼1200 F2 progeny were then plated on 1 × 10-cm RNAi plates. 45 hours after plating, animals were washed with M9 into a 15 ml centrifuge tube and spun at 3,000 rpm for 30 s. Supernatant was aspirated, and worms were then transferred to a 1.5 ml microcentrifuge tube. All samples were washed with M9 and supernatant aspirated down to 100 μl and 1 ml M9 was added then spun down on minifuge for 20 seconds. This wash step was performed three times. Then, 100 μl of room temperature 10x OP50, 146 μl of M9 and 3.9 μl of *N. parisii* spores (∼20 × 10^6 spores) were mixed with each sample and plated onto pre-dried 6-cm unseeded NGM plates and spread evenly across the plate. Plates were left to dry in laminar flow hood for 30 minutes, then moved to 25 °C for 3 h. All samples were then washed off 6-cm plates, washed with M9 three times, and plated onto fresh 10-cm RNAi plates, which were then incubated at 25 °C for an additional 27 h. Worms were then fixed in acetone and stained with FISH probes conjugated to the red Cal Fluor 610 fluorophore that hybridize to *N*. *parisii* ribosomal RNA^49^ (Biosearch Technologies, Hoddeson, United Kingdom) and incubated at 47 °C overnight (16-18 h). Animals were then washed and imaged using the ImageXpress Nano plate reader using the 4x objective (Molecular Devices, LLC, San Jose, CA). Worms were automatically segmented using the protocol/code located at GitHub (https://github.com/TroemelLabUCSD/Mitochondria-and-PathogenLoad-Analysis) using MetaXpress 2018 software version 6.5.2.351.

### Statistics

Statistical analysis was performed using Prism 9 & 10 software (GraphPad). Before statistical analysis, the D’Agostino & Pearson and Shapiro-Wilk tests were performed for all experiments to assess the distribution of the data. If data were normally distributed, standard parametric statistical tests were performed. Nonparametric tests were performed for non-normally distributed data, as determined by either the D’Agostino & Pearson or the Shapiro-Wilk tests. The corresponding figure legends describe the statistical test for each experiment, the number of data points analyzed, and the number of experimental replicates performed.

## Acknowledgements

We are grateful to Noelle Antao, Michalis Barkoulas and Vladimir Lazetic for comments on the manuscript. We thank M. Wood at Scripps Research Institute for assistance in TEM experiments. We thank Gira Bhabha and Damian Ekiert for helpful conversations about PALS protein structure. Some strains were provided by the CGC, which is funded by NIH Office of Research Infrastructure Programs (P40 OD010440). This work was supported by NIH under R01 AG052622, GM114139, AI176639 and by NSF IOS-2301657 to E.R.T; S.S.G. received funding from NIH award K12GM068524. M.W.S. received funding from NIH award GM007240.

## Figure Legends

**Figure S1.**
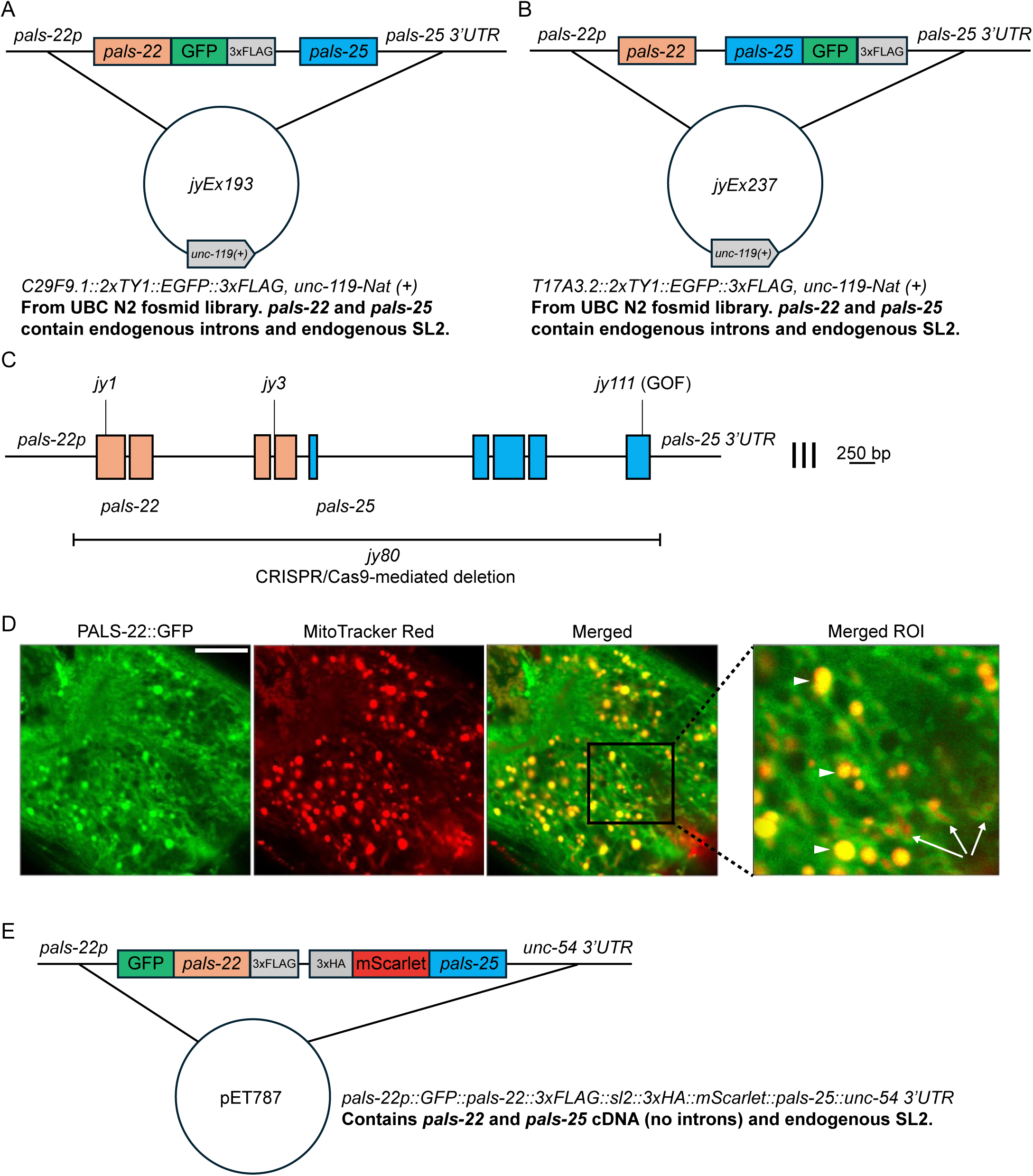
PALS-22 and PALS-25 expression vectors used in this study, and intestinal localization of PALS-22::GFP. **A-B)** Simplified vector maps of the TransgeneOme fosmid constructs used to express PALS-22::GFP and PALS-25::GFP. **A)** *jyEx193[pals-22p::pals-22::GFP::3xFLAG::sl2::pals-25, unc-119(+)]* expresses PALS-22 C-terminally tagged with GFP::3xFLAG and wild-type PALS-25 from the same fosmid using the endogenous *pals-22/25* regulatory elements. **B)** *jyEx237[pals-22p::pals-22::sl2::pals-25::GFP::3xFLAG, unc-119(+)]* expresses PALS-25 C-terminally tagged with GFP::3xFLAG and wild-type PALS-22 from the same fosmid using the endogenous *pals-22/25* regulatory elements. **C)** The *pals-22* and *pals-25* genomic locus. *pals-22* (orange exons) and *pals-25* (blue exons) are in an operon. **D)** PALS-22::GFP colocalizes with the mitochondria marker MitoTracker Red in the intestine. The first ring of intestinal cells of an adult animal is shown. White arrows denote PALS-22::GFP localized to mitochondria, white triangles denote autofluorescent gut granules. Scale bar = 10 μm. **E**) Vector map of pET787 used to co-express fluorescently tagged GFP::PALS-22 and mScarlet::PALS-25 using a smaller DNA construct lacking endogenous introns but maintaining the endogenous promoter and SL2 sequence for *pals-22/25*.

**Figure S2.**
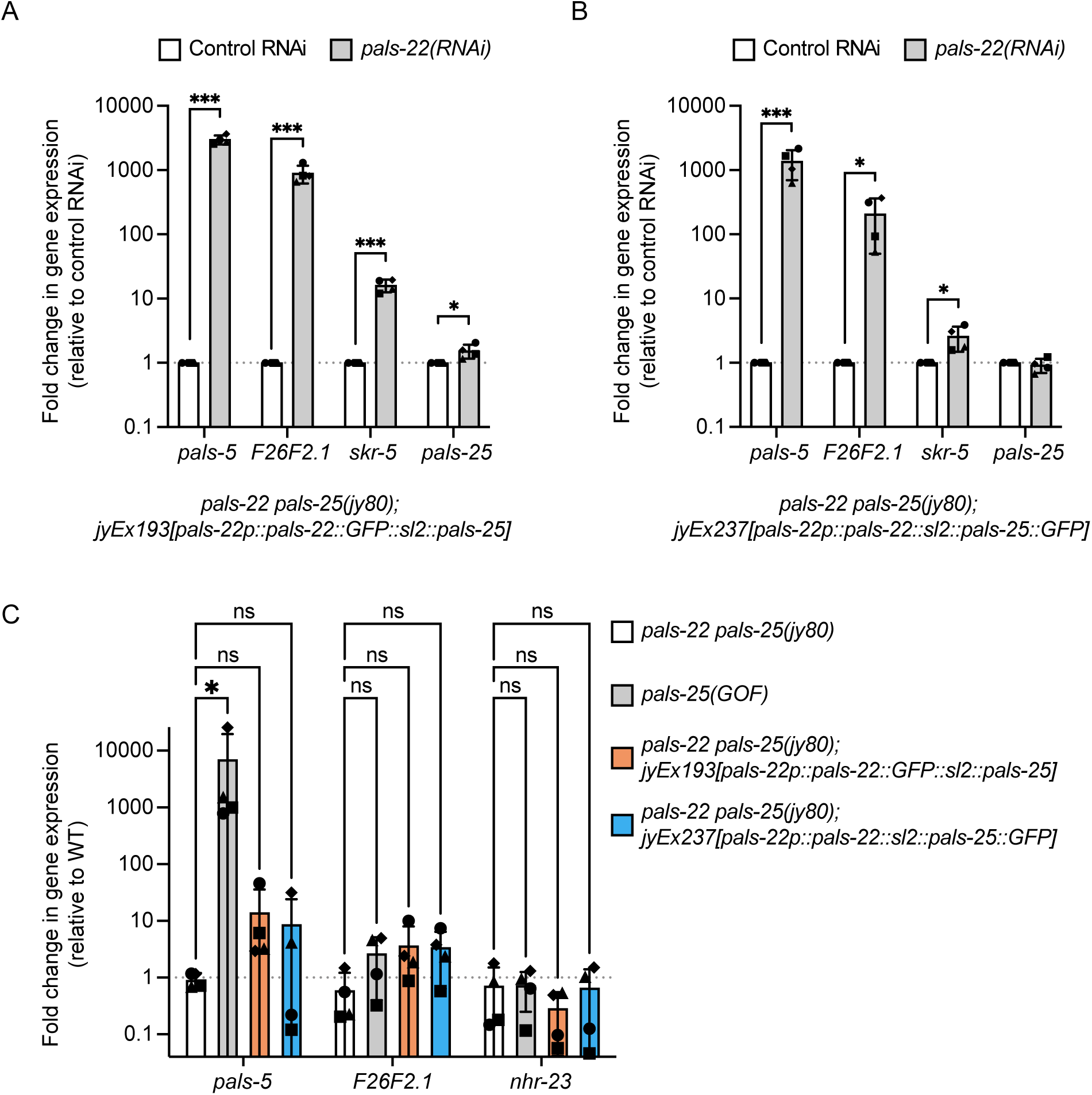
PALS-22::GFP and PALS-25::GFP are functional for regulating the IPR. **A)** RT-qPCR of select IPR-induced genes (*pals-5*, *F26F2.1*, and *skr-5)* in *pals-22 pals-25(jy80)* double mutants expressing *jyEx193* following either control RNAi or *pals-22* RNAi. IPR gene expression is induced following *pals-22* RNAi, indicating that PALS-22::GFP expressed from *jyEx193* is functional for the repression of PALS-25. **B)** RT-qPCR of select IPR-induced genes (*pals-5*, *F26F2.1*, and *skr-5)* in *pals-22 pals-25(jy80)* double mutants expressing *jyEx237* following either control RNAi or *pals-22* RNAi. IPR gene expression is induced following *pals-22* RNAi, indicating that PALS-25::GFP expressed from *jyEx237* is functional for activating the IPR. For **A** and **B**, although *pals-22* and *pals-25* are co-transcribed in an operon, *pals-22* RNAi does not knock down *pals-25* transcript levels. *\* p <* 0.05, *\*\*\* p <* 0.001, One-tailed t-test. **C)** RT-qPCR of select IPR-induced genes *pals-5*, *F26F2.1,* and control gene *nhr-23,* in *pals-22 pals-25(jy80)* double mutants in white, *pals-25(GOF)* in gray, *pals-22 pals-25(jy80)* double mutants expressing *jyEx193* in orange, and *pals-22 pals-25(jy80)* double mutants expressing *jyEx237* in blue. *pals-5* IPR gene expression is induced in *pals-25(GOF)* relative to *pals-22/25* double mutants but not in *pals-22/25* double mutants expressing *jyEx193* nor in *pals-22/25* double mutants expressing *jyEx237*, indicating that the IPR is not induced in basal conditions with these TransgeneOme fosmid constructs. *pals-25(GOF)* does not induce *F26F2.1,* which is known not to be induced in this mutant (Gang et al. 2022). *\* p <* 0.05, two-way ANOVA. For **A-C**, n = 4 independent experimental replicates, different symbol shapes represent the expression values for replicates performed on different days. The graphs indicate mean values and error bars represent standard deviations.

**Figure S3.**
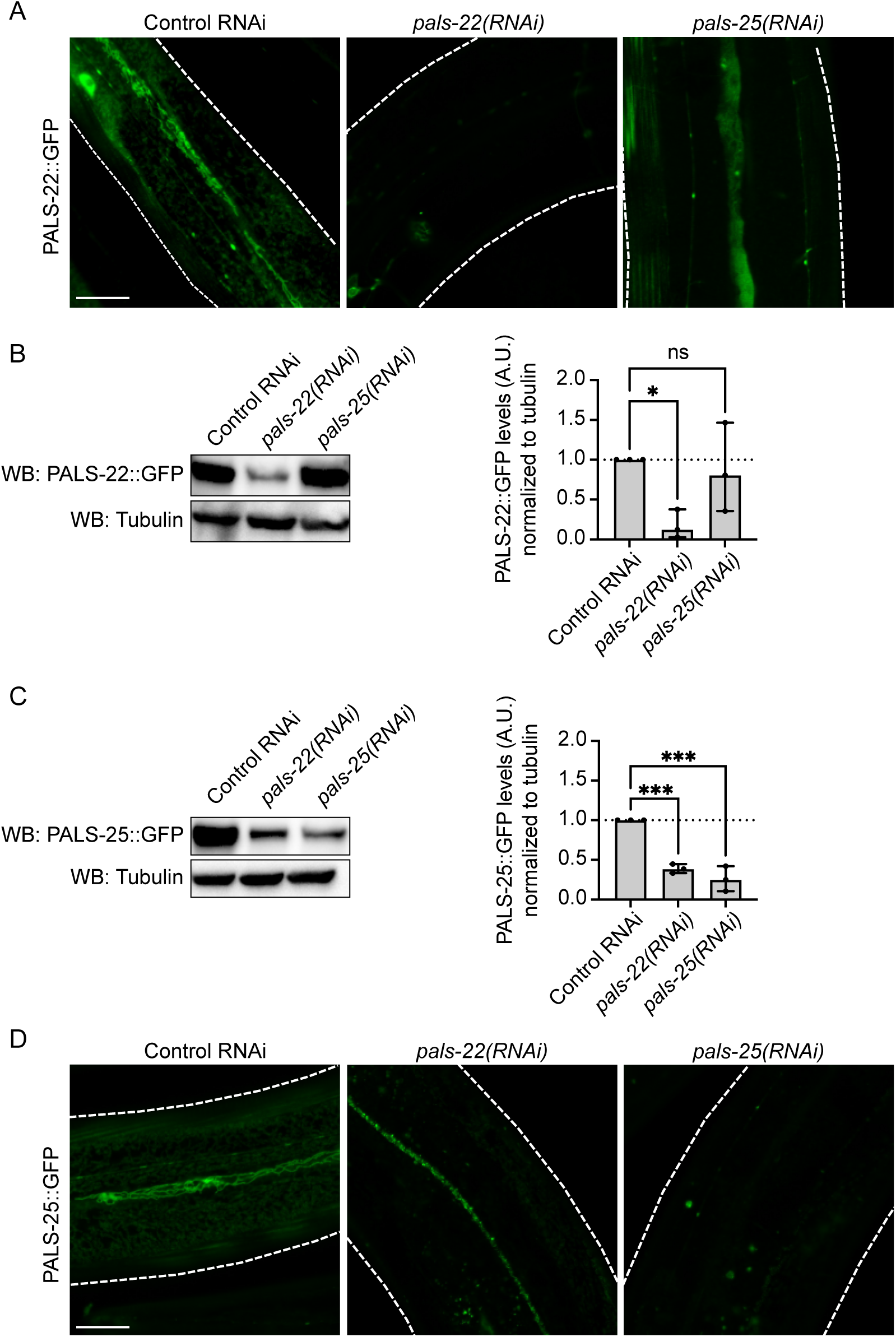
PALS-22 protein levels do not change upon knock-down of *pals-25,* but PALS-25 protein levels are reduced upon knock-down of *pals-22*. **A)** Representative images of PALS-22::GFP expressed in the epidermis of adult animals under control, *pals-*22, and *pals-*25 RNAi treatments. **B)** Left, Western blot analysis of PALS-22::GFP expression levels in total protein lysates following different RNAi treatments using custom antibodies against PALS-22 and commercially available antibodies against tubulin as a loading control. Right, Quantification of PALS-22::GFP levels in total protein lysates following the different RNAi treatments, normalized to tubulin. **C)** Left, Western blot analysis of PALS-25::GFP expression levels in total protein lysates following different RNAi treatments using custom antibodies against PALS-25 and commercially available antibodies against tubulin as a loading control. Right, Quantification of PALS-25::GFP levels in total protein lysates following the different RNAi treatments, normalized to tubulin. **D)** Representative images of PALS-25::GFP expressed in the epidermis of adult animals under control, *pals-*22, and *pals-*25 RNAi treatments. For **A** and **D**, scale bar = 20 μm. For **B** and **C**, * *p <* 0.05, *** *p <* 0.001, one-way ANOVA with Dunnett’s multiple comparisons test compared to control RNAi-treated animals. n = 3 independent experimental replicates. The graphs indicate mean values and error bars represent standard deviations.

**Figure S4.**
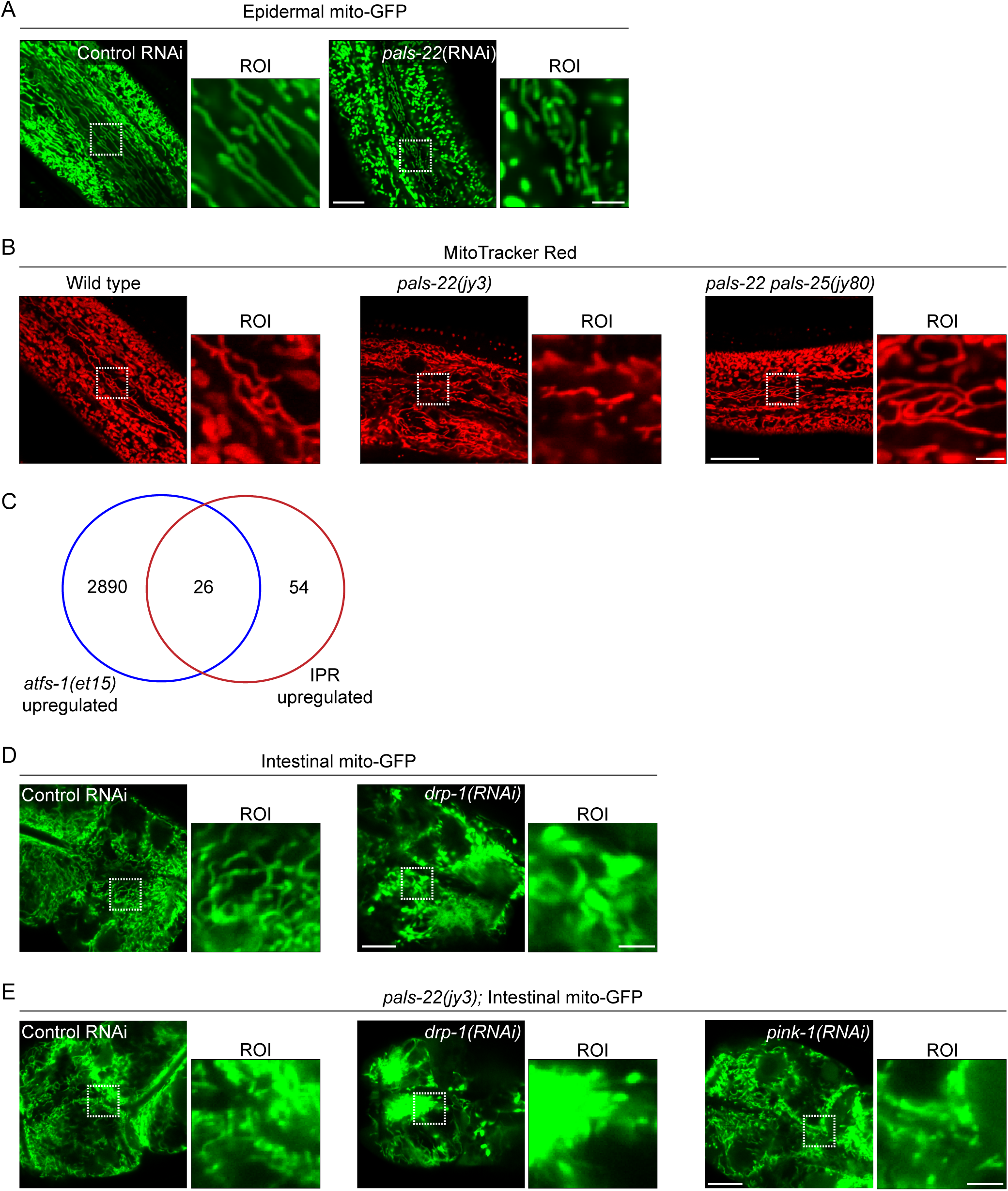
Images of mitochondria morphology upon loss of *pals-22* and/or *pals-25*. **A)** Representative images of *jyEx4796[col-19p::*mito-GFP] expression in the epidermis of wild-type adult animals treated with control or *pals-22* RNAi; *pals-22* RNAi alters mitochondrial morphology in the epidermis. **B)** Representative images of wild-type, *pals-22(jy3)*, and *pals-22 pals-25(jy80)* mutants stained with MitoTracker Red to visualize epidermal mitochondria morphology; *pals-22 pals-25(jy80)* mutants display grossly wild-type mitochondria morphology. Scale bar = 20 μm, ROI scale bar = 2.5 μm. **C)** mitoUPR upregulated genes constitutively induced by the *atfs-1(et15)* gain-of-function allele have significant overlap with the IPR. Hypergeometric test, RF = 2.2, *p* < 4.036e-05. **D)** Representative images of *mgIs48[ges-1p::mito-GFP]* animals treated with control or *drp-1* RNAi. *drp-1* RNAi alters mitochondrial morphology in the intestine. **E)** Representative images of *pals-22(jy3); mgIs48[ges-1p::mito-GFP]* animals treated with control, *drp-1,* or *pink-1* RNAi. *drp-1* and *pink-1* RNAi do not rescue mitochondrial morphology defects observed in *pals-22* mutant animals. For **A, D,** and **E**, scale bar = 10 μm, ROI scale bar = 2.5 μm.

**Figure S5.**
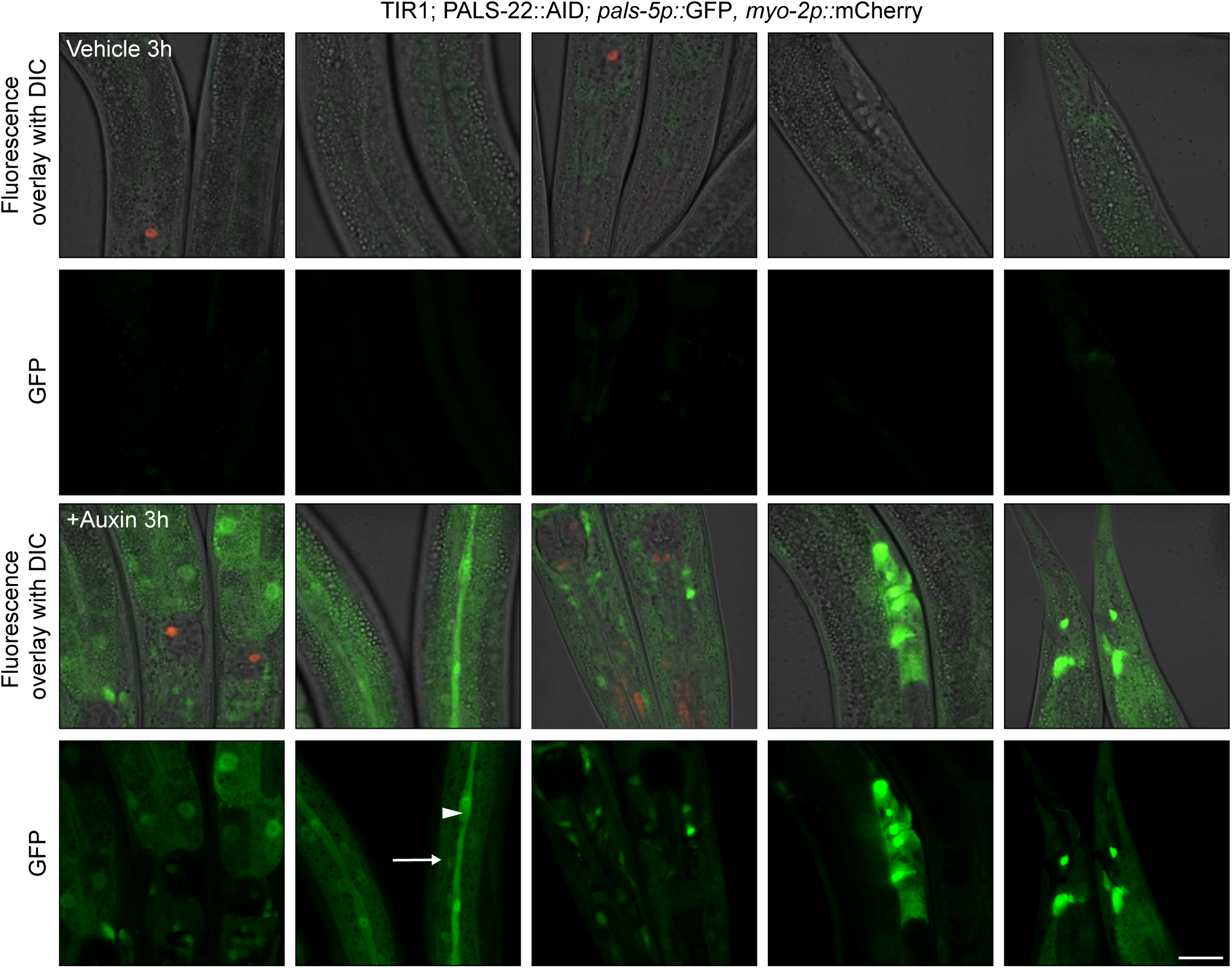
IPR reporter expression is induced in multiple tissues following 3 hours of auxin-mediated depletion of PALS-22. IPR reporter expression in transgenic animals expressing ubiquitous TIR1, endogenous *pals-22* tagged with AID, and the *pals-5p::*GFP, *myo-2p::*mCherry IPR reporter treated with vehicle control (top two rows) or auxin (bottom two rows) for 3 h starting at the L4 life stage. Top images for each treatment are an overlay of DIC, GFP, and mCherry fluorescence channels. Bottom images are GFP alone and indicate the induction of the *pals-5p::*GFP IPR reporter. IPR expression is visible after 3 h auxin treatment in multiple tissues; left to right = the anterior intestine, the epidermis (seam cells denoted by the white triangle; other epidermal cells denoted by white arrow), amphid, pre-vulva, posterior intestine and neurons near the rectum. Scale bar = 20 μm.

**Figure S6.**
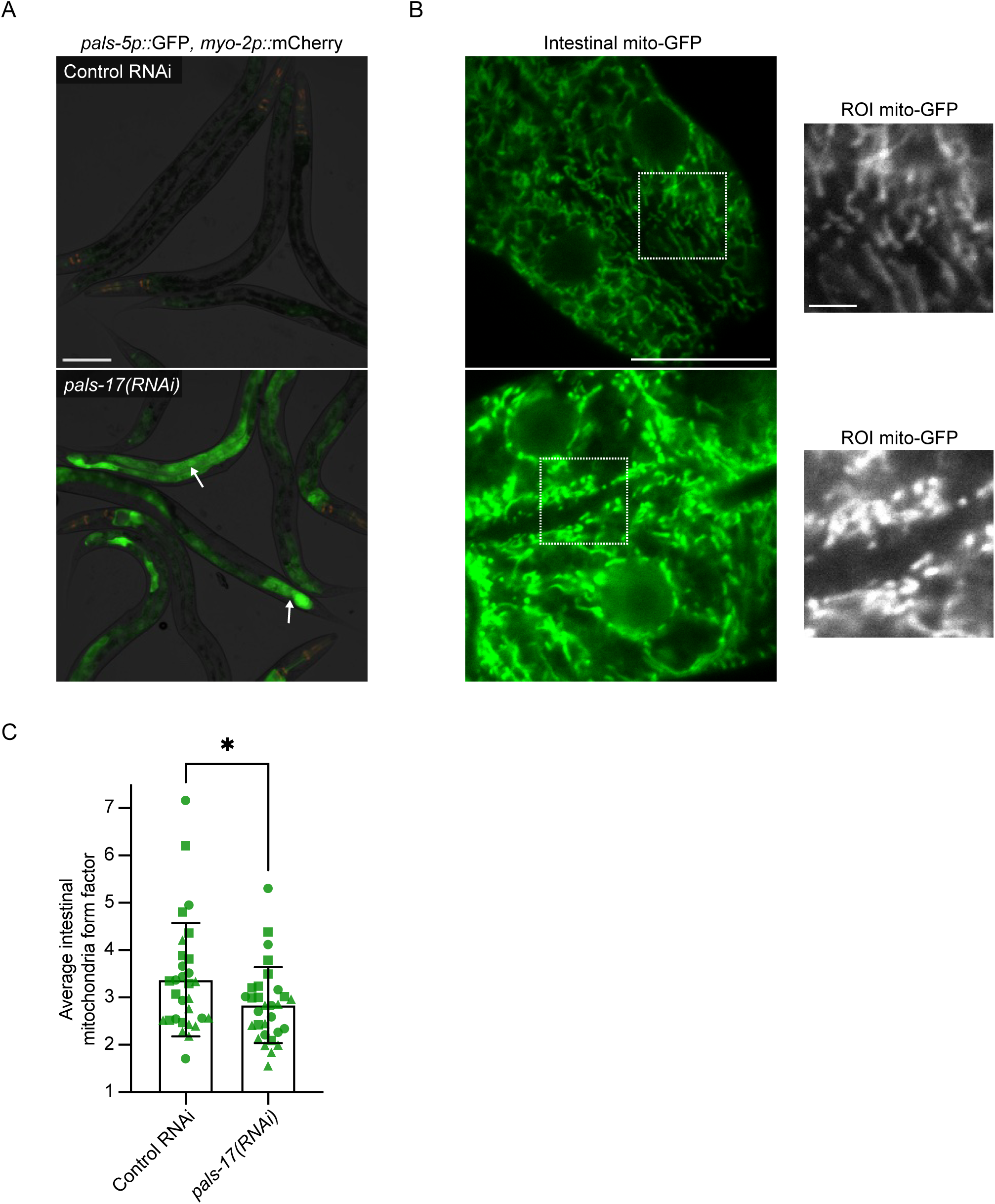
Knock-down of *pals-17* induces the IPR and changes intestinal mitochondria morphology. **A)** Representative images of *pals-5p::*GFP, *myo-2p::*mCherry IPR reporter expressing animals treated with either control or *pals-17* RNAi for 24 h starting at 20 h post-L1. Control RNAi does not induce IPR reporter, and *pals-17* RNAi induces *pals-5p*::GFP IPR reporter expression in the intestine at the 24 h timepoint shown (white arrows). *myo-2p::*mCherry is part of the same transgene and is constitutively expressed in the pharynx of transgenic animals at all life stages. Images are an overlay of DIC, GFP, and mCherry fluorescence channels. Scale bar = 100 μm. **B)** Representative images of *mgIs48[ges-1p::mito-GFP*] animals 20 h post-L1 exposed to control or *pals-17* RNAi for 24 hours. The white boxes indicate an ROI in the first ring of intestinal cells, and increased magnification showing mitochondria morphology in the ROI based on mito-GFP expression is inset. Scale bar = 15 μm, ROI scale bar = 2.5 μm. **C)** *pals-17* RNAi induces changes in intestinal mitochondria morphology, as measured by form factor. *\* p <* 0.05, unpaired t-test. n = 30 ROIs analyzed per condition, two ROIs per animal, across three experimental replicates. The graph indicates mean values and error bars represent standard deviations. Different symbol shapes represent ROIs from animals imaged on different days.

